# HSF1 remodels mitochondrial biogenesis and function in cancer cells via TIMM17A

**DOI:** 10.1101/2025.05.12.653547

**Authors:** Ngoc G.T. Nguyen, Hem Sapkota, Yoko Shibata, Aleksandra Fesiuk, Matthew Antalek, Vibhavari Sail, Daniel J. Ansel, David R. Amici, Frederick F. Peelor, Holly Van Remmen, Benjamin F. Miller, Marc L. Mendillo, Richard I. Morimoto, Jian Li

## Abstract

Mitochondria play critical roles in energy production and cellular metabolism. Despite the Warburg effect, mitochondria are crucial for the survival and proliferation of cancer cells. Heat Shock Factor 1 (HSF1), a key transcription factor in the cellular heat shock response, promotes malignancy and metastasis when aberrantly activated. To understand the multifaceted roles of HSF1 in cancer, we performed a genome-wide CRISPR screen to identify epistatic interactors of HSF1 in cancer cell proliferation. The verified interactors of HSF1 include those involved in DNA replication and repair, transcriptional and post-transcriptional gene expression, and mitochondrial functions. Specifically, we found that HSF1 promotes cell proliferation, mitochondrial biogenesis, respiration, and ATP production in a manner dependent on TIMM17A, a subunit of the inner membrane translocase. HSF1 upregulates the steady-state level of the short-lived TIMM17A protein via its direct target genes, HSPD1 and HSPE1, which encode subunits of the mitochondrial chaperonin complex and are responsible for protein refolding once imported into the matrix. The HSF1- HSPD1/HSPE1-TIMM17A axis remodels the mitochondrial proteome to promote mitochondrial translation and energy production, thereby supporting robust cell proliferation. Our work reveals a mechanism by which mitochondria adjust protein uptake according to the folding capacity in the matrix by altering TIM complex composition.

## Introduction

Heat Shock Factor 1 (HSF1) is best known for its role in the cellular heat shock response (HSR), during which HSF1 induces the transcription of molecular chaperones and other proteostatic genes to manage misfolded proteins caused by elevated temperature ^1^. HSF1 also plays essential roles in animal development, reproduction, and longevity ^2^, and supports malignancy ^3,4^ through transcriptional programs that differ from the HSR ^5–7^. The activation of HSF1 in cancer cells, indicated by nuclear HSF1 levels and the expression of its target genes, is associated with poor outcomes in patients ^7,8^. A potent HSF1 inhibitor that directly binds to HSF1 and stimulates its degradation significantly inhibits cell proliferation and tumor progression in prostate cancers ^9^.

Despite the strong correlation with clinical outcomes and therapeutic potential, the mechanistic understanding of HSF1’s roles in malignancy remains incomplete. HSF1 has been reported to promote cancer cell proliferation, survival, and metastasis by regulating proteome stability ^10,11^, DNA repair ^12^, mRNA processing and protein synthesis ^13,14^, energy metabolism ^15–17^, and cell migration ^18^. Evaluating the relative contributions of HSF1’s diverse functions in a specific cancer model is lacking and remains challenging. These processes are proposed to be either mediated by HSF1 target genes or independent of HSF1’s transcriptional activities. Since HSF1’s transcriptional targets and protein-protein interactions tend to be context-specific ^7,19^, it is difficult to predict whether the reported mechanisms from one cancer type apply to another.

To comprehensively understand the cellular pathways through which HSF1 promotes cancer cell proliferation and link them to the HSF1 transcriptional program, we combined an epistatic interactor screen with multi-omics analyses for HSF1 genomic binding and its impacts on the transcriptome and proteome. Our functional genomics study rediscovered the reported functions of HSF1 and pointed to several previously unappreciated pathways that underlie HSF1’s contribution to rapid cell proliferation. Specifically, we uncovered that HSF1 remodels mitochondria and energy metabolism by regulating the inner membrane translocase.

## Results

### Epistatic interactors of HSF1 play critical roles in DNA replication and repair, gene expression, and mitochondrial functions

To dissect the multifaceted functions of HSF1 in cancer, we performed paired CRISPR knockout screens in cells with either wild-type HSF1 or HSF1-null to determine the epistatic interactors of HSF1 in cell proliferation (Fig. 1A&B). We chose the aggressive prostate cancer cell line PC3M as our model, which has been shown to depend on HSF1 for rapid proliferation in cell culture and tumor progression in xenograft models ^20^. Loss of HSF1 increased the doubling time by approximately 12% (Fig. S1A), which is sufficient to resolve genetic buffering effects during negative selection while allowing us to detect synthetic lethal interactions. Based on the types of genetic interactions (additive, genetic buffering, or synthetic lethal), we could position HSF1 within established cellular pathways (Fig. 1A) and gain a comprehensive understanding of its roles in aberrant cancer cell proliferation. We fitted the time-dependent changes of sgRNAs in the Brunello library (four sgRNAs per gene) to an exponential decay model to calculate the CRISPR score (the beta value, Fig. 1C & S1B), which measures the impacts of a given gene knockout on cell proliferation and survival. We then compared the CRISPR scores from the parental (PC3M- WT) and HSF1 knockout cells (HSF1-KO) using the following criteria to determine potential epistatic interactions: 1> the delta CRISPR score (PC3M-WT minus HSF1-KO) of a given gene (average of four sgRNAs per gene) is significantly different from the average of the non-target control pool (Fig. S1C, P<0.025). Our sequencing data recovered 994 out of 1000 control sgRNAs in the Brunello library, the delta CRISPR scores of which are almost evenly distributed around zero; 2> the paired t-test of CRISPR scores for the four individual sgRNAs of a given gene indicates a significant difference between PC3M-WT and HSF1-KO (FDR: 0.05). This filters out false positives caused by outlier behavior from one or two sgRNAs; 3> the potential genetic interactors were identified in both pairs of screens that used different HSF1-KO clones (Fig. 1D). Using these stringent criteria, we identified 91 genetic buffering interactors and 24 synthetic lethal interactors of HSF1.

**Fig. 1.**
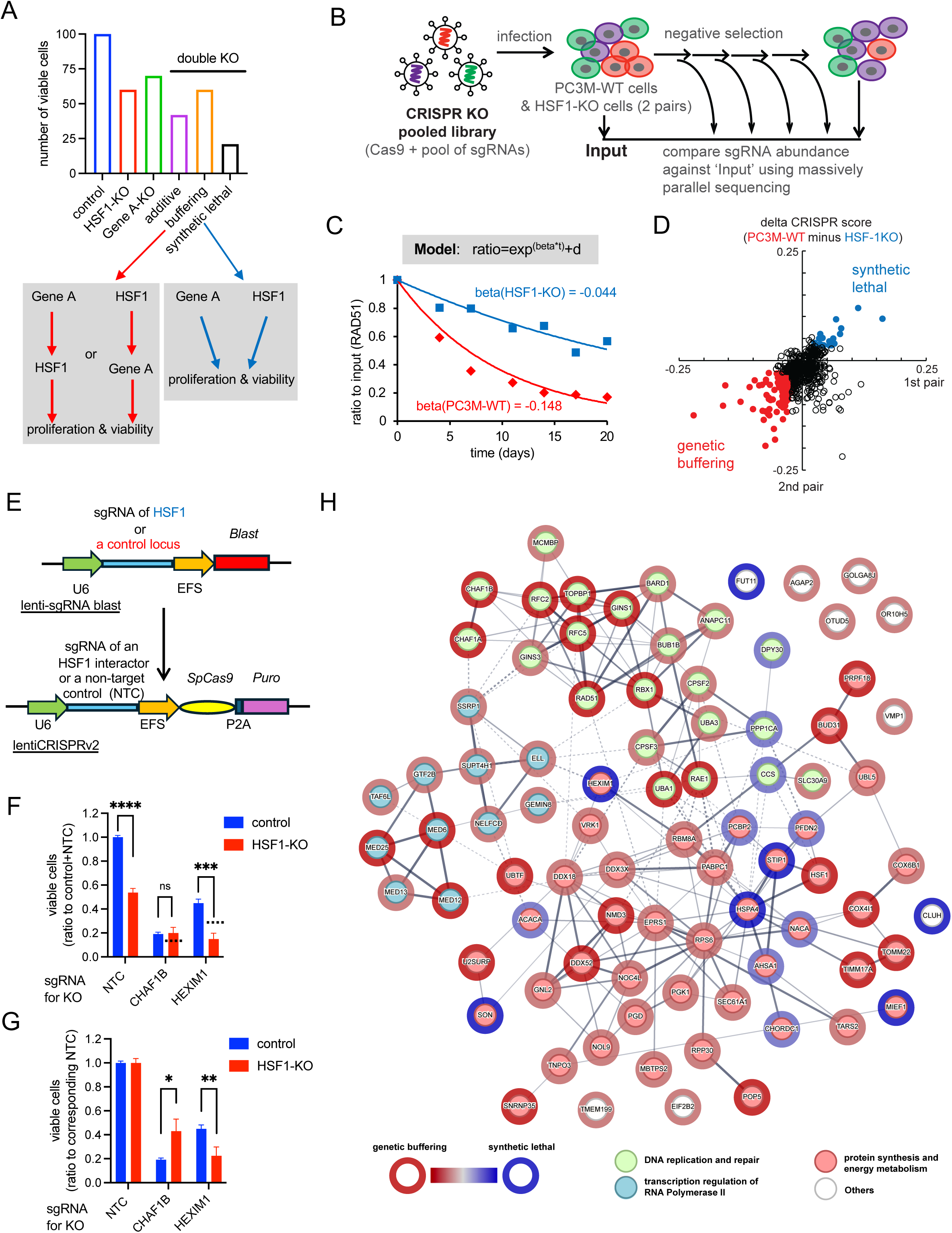
HSF1 interactors are involved in critical cellular processes. (A) Schematic diagram of epistatic interactions by comparing cell proliferation defects caused by single and double knockout (KO) of HSF1 and another essential gene ‘A’. The ratio (%) of viable cells compared to that in the control is plotted in the histograms. The additive effect: ratio (double KO) = ratio (HSF1-KO) * ratio (Gene A-KO). (B) Experimental strategy: A population of PC3M wild-type cells (PC3M-WT) or HSF1-KO cells was infected with a pooled sgRNA library and collected at different time points (every 3 to 4 days) during negative selection. The abundance of sgRNA-encoding constructs was determined by deep sequencing and compared to the input samples at Day 0. Two clones of PC3M-WT and HSF1-KO cells were used in the paired screens. (C) CRISPR score calculation using RAD51 as an example. The relative abundance of sgRNA over time was fitted into an exponential decay model. The beta value, representing how quickly the RAD51-KO cells declined in the cell population, is calculated as the CRISPR score. RAD51 sgRNA decreased more rapidly in the PC3M-WT cells than in the HSF1-KO cells. Correspondingly, RAD51 exhibits a smaller CRISPR score in PC3M-WT cells than in HSF1-KO cells. Dots on the graph represent the weighted average abundance of the four sgRNAs targeting RAD51 at each time point, which are utilized for the beta value calculation. (D) Scatter plots illustrate the delta CRISPR scores (PC3M-WT minus HSF1-KO) for genes essential in either cell type. Blue and red dots represent the overlapped synthetic lethal and genetic buffering interactors of HSF1, respectively, in both paired screens. (E) Schematic diagram of the co-CRISPR assay. The sgRNA targeting the MAPT1 gene or the AAVS locus serves as the control for HSF1 sgRNA during the first infection. The sgRNA of an HSF1 interactor or a non-target control (NTC) was co-expressed with Cas9 during the second infection. (F & G) Histograms showing the number of viable cells measured by PrestoBlue assays on Day 7. The ratios to the control + NTC cell numbers (F) and the ratios to the corresponding NTC in either the control or HSF1-KO cells (G) are presented (mean ± SD, n=3). Dashed lines represent the calculated additive effects. Unpaired t-test: ns P >= 0.05; * P < 0.05; ** P < 0.01; *** P < 0.001; **** P < 0.0001. (H) Network of verified epistatic interactors of HSF1. The physical and functional interaction network was retrieved from the STRING database and grouped using k-means clustering (n=3). The inner circle color of each node represents the cluster to which the gene belongs. The halo color of each node indicates the type of genetic interaction (red: genetic buffering; blue: synthetic lethal), while the darkness reflects the confidence (verified in one of the co-CRISPR experiments or both). The color saturation of edges denotes the confidence score of a functional interaction between two interactors.

We then tested the interactors individually in double knock-out cells (interactor + HSF1). Since HSF1 may play a role in viral infections ^21^, we designed the assay (Fig. 1E-G) to control potential differences caused by HSF1 knockout in cellular responses to lentiviral infection. First, we introduced an sgRNA targeting HSF1 or a control locus without expressing the Cas9 protein. After that, we infected the cells with lentivirus co-expressing Cas9 and an sgRNA for either one candidate interactor or a non-target control (NTC). The second infection resulted in the KO of HSF1, the candidate interactor, or the simultaneous KO of both. We included an sgRNA targeting a control locus to ensure that CRISPR-induced double-strand DNA breaks also occur in the negative control cells (the control locus + NTC). This is particularly helpful because several candidate interactors of HSF1 are involved in the DNA damage response (Table S1). Our study used the MAPT gene, a lowly expressed, non-essential gene in PC3M and a previously used control ^22^, or the AAVS locus, known as a ‘safe harbor’ for transgenes ^23^, as the control locus. We conducted independent co-CRISPR assays in two laboratories using different sets of sgRNAs, and more than two-thirds of the candidate interactors (67 of 91 genetic buffering interactors and 16 of 24 synthetic lethal interactors) were confirmed by at least one co-CRISPR validation (Fig. 1F-H, S1 D-I).

The verified epistatic interactors of HSF1 form a highly connected network enriched with physical and genetic interactions (Fig. 1H). Many have well-established roles in rapid cell proliferation, encompassing DNA replication and repair, transcriptional and post-transcriptional gene expression, and energy metabolism, implicating HSF1’s involvement in these critical cellular processes. The knockout of components in DNA replication and repair pathways exhibited genetic buffering effects in the HSF1 null cells. These include two subunits of the GINS complex (GINS1 and GINS3) that unwind DNA for the establishment and progression of the replication fork ^24^, along with two subunits of Replication Factor C (RFC2 and RFC5) that are essential for both DNA replication and repair ^25^, as well as TOPBP1 and RAD51, which collaborate in homologous recombination (HR) to repair damaged DNA ^26^. This result supports a role of HSF1 in DNA replication and/or damage repair. Similarly, several critical regulators in RNA polymerase II (Pol II) transcription displayed genetic buffering effects with HSF1, including four subunits of the Mediator complex and subunits of multiple transcription elongation factors such as NELF (NELFCD), DSIF (SUPT4H1), FACT (SSRP1), and SEC (ELL). HSF1 has been shown to physically or functionally interact with these transcription factors for gene regulation in the HSR ^27–29^. HSF1 may utilize similar transcription machinery to drive its cancer-specific transcription program or affect the activities of these transcription factors. HSF1 also genetically interacts with a large group of genes that function at different steps of protein synthesis. Loss of HSF1 buffers the detrimental effects from the knockout of components in rRNA and tRNA processing (NOL9, POP5, and RPP30) ^30–32^, ribosome assembly and export (NOC4L and NMD) ^33,34^, and tRNA aminoacylation (EPRS1)^35^, supporting the idea that HSF1-dependent chaperone expression facilitates robust protein translation. Conversely, HSF1 knockout is synthetic lethal upon loss of any of the following chaperone and co-chaperone genes (NACA, PFDN2, HSPA4, STIP1, AHSA1, and CHORDC1), consistent with proteostasis being on the verge of collapse in HSF1- KO cells, which cannot afford further reduction in protein folding capacity. Finally, seven mitochondrial genes show epistatic interactions with HSF1. They are involved in mitochondrial protein transport (TIMM17A and TOMM22), electron transport chain function (COX4L1 and COX6B1), regulation of mitochondrial morphology (MIEF1 and CLUH), and mitochondrial translation (TARS2), suggesting that HSF1 may influence multiple aspects of mitochondrial activities.

### HSF1 remodels the steady-state mitochondrial proteome

HSF1 is a well-established transcription regulator of stress responses and cancer cell proliferation, although through distinct transcriptional programs ^7^. We wonder how many of HSF1’s genetic interactions are linked to its roles in gene expression. We performed both RNA-seq and proteomics analyses to test whether any of the confirmed HSF1 interactors significantly altered protein levels and, if so, whether these changes resulted from mRNA alterations. If the genetic buffering or synthetic lethality effects of HSF1 knockout occur through regulating the expression of its interactors, the protein levels of these interactors are expected to decrease for genetic buffering and increase for synthetic lethality. 23 out of 83 HSF1 genetic interactors changed protein levels by at least 25% (FDR: 0.05) upon HSF1 knockout (Fig. 2A). However, most of these changes do not support the model that HSF1-dependent gene expression directly dictates the genetic interaction. 5 of 6 synthetic lethal interactors decreased protein levels in HSF1-KO cells, 4 of which encode co-chaperones of HSP70 and HSP90 (STIP1, AHSA1, HSPA4, and CHORDC1). Published work ^20^ and our data indicate that HSF1 directly activates the transcription of members of HSP70 and HSP90 chaperones in PC3M cells (Fig. 5C). Therefore, when HSP70 and HSP90 protein levels are low (Table S2), the loss of co-chaperones that promote their substrate binding or folding activities is synthetic lethal. Conversely, 12 of 17 genetic buffering interactors increase protein levels in the HSF1-KO cells. Notable exceptions are the 4 proteins involved in energy metabolism, of which the two enzymes involved in sugar metabolism, PGD and PGK1, decreased both protein and mRNA levels in HSF1-KO cells, and the two mitochondrial proteins, TIMM17A and TARS2, decreased protein but not mRNA levels in the absence of HSF1. Our analyses suggest that the transcription regulation of its epistatic interactors by HSF1 is not the major underlying mechanism for the identified genetic interactions, which likely involve additional partners and regulatory mechanisms that impact the activities of HSF1 interactors.

**Fig. 2.**
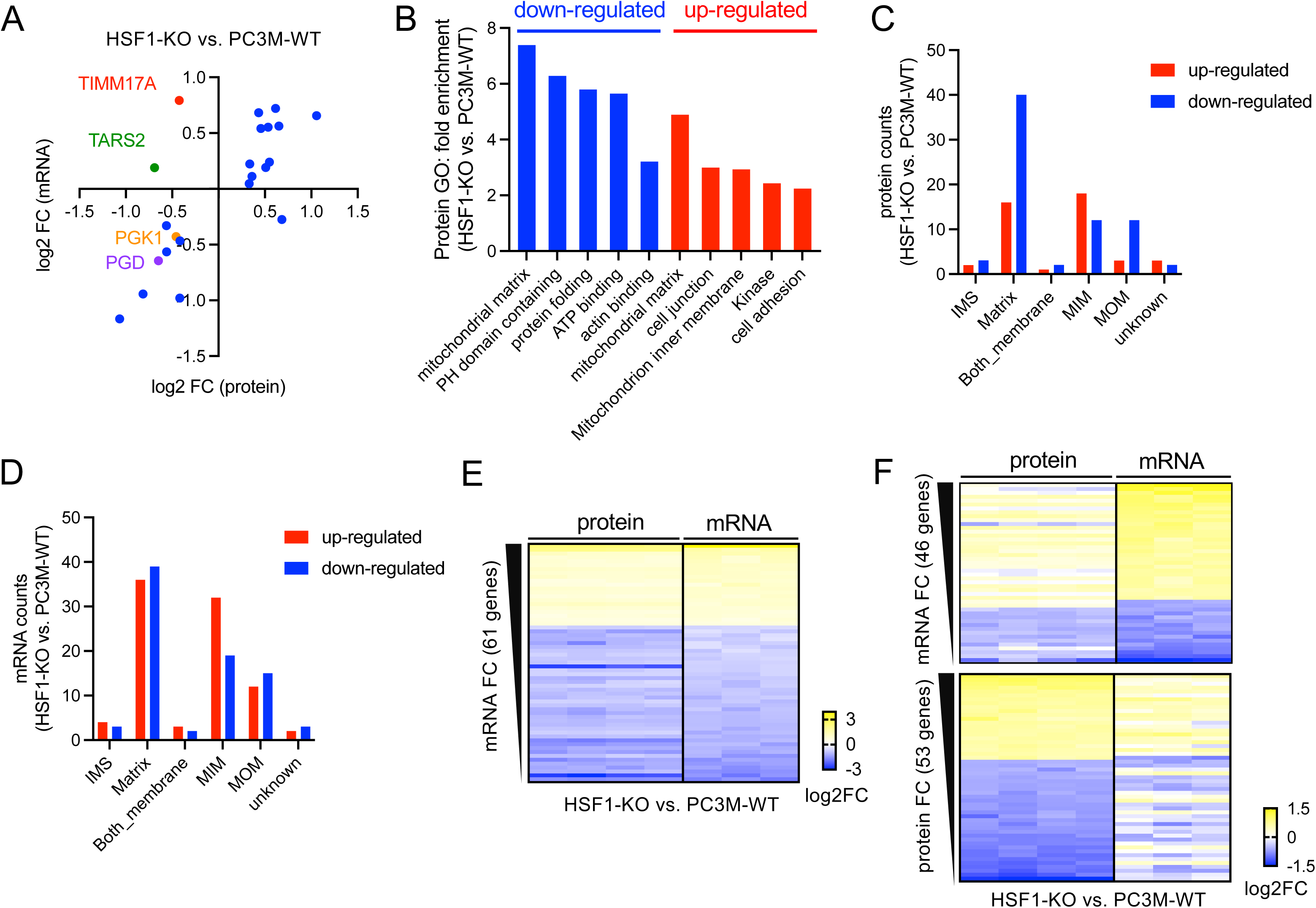
HSF1 remodels the steady-state mitochondrial proteome. (B) Scatter plots displaying the log2 fold change (FC) of protein (x axis) and mRNA (y axis) of HSF1 epistatic interactors following the knockout of HSF1 in PC3M cells. The epistatic interactors of HSF1 that altered protein levels by at least 1.25-fold (FDR: 0.05) are included. RNA-seq was performed in biological triplicate, and TMT-proteomics was performed in four biological replicates. The average fold changes (HSF1-KO vs. PC3M-WT) in mRNA and protein levels are plotted. (C) Histograms showing gene ontology (GO) analysis of differentially expressed proteins following the knockout of HSF1 in PC3M cells (1.5-fold, FDR: 0.05). (C&D) Histograms displaying the numbers of differentially expressed proteins following HSF1 knockout in PC3M cells across various mitochondrial compartments. Mitochondrial genes that significantly altered their protein (C) or mRNA (D) levels (1.5-fold, FDR: 0.05) were included. IMS: intermembrane space; MIM: mitochondrial inner membrane; MOM: mitochondrial outer membrane. (E&F) Heatmap showing log2 fold change (FC) of protein and mRNA of differentially expressed mitochondrial genes upon knockout of HSF1 in PC3M cells. 61 genes significantly changed both their mRNA and protein levels (1.5-fold, FDR: 0.05) (E). 99 genes significantly changed either their mRNA (F, upper panel) or protein levels (F, lower panel) but not both.

We then extended the proteomics analyses to a genome-wide scale and found that HSF1 significantly impacts the mitochondrial proteome (Fig. 2B). Upon HSF1 knockout, both down- and up-regulated proteins (1.5-fold, FDR: 0.05) are most enriched with those residing in the mitochondrial matrix. Additionally, mitochondrial inner membrane proteins are also enriched in the up-regulated group. This result indicates that the steady-state mitochondrial proteome is dramatically remodeled following the loss of HSF1. By cross-comparing the mRNA and protein data, we found that, although there are similar numbers of down- and up-regulated mitochondrial genes at the mRNA level (up: 89; down: 81, 1.5-fold, FDR: 0.05), a much higher number of mitochondrial proteins are down-regulated (up: 43; down: 70), especially those in the matrix (Fig. 2C & D). Indeed, among the 160 mitochondrial genes differentially expressed at the level of mRNA or protein, only 61 genes (∼38%) significantly changed in the same direction at both mRNA and protein levels (Fig. 2E & F), suggesting that post-transcriptional mechanisms play an essential role in HSF1-mediated mitochondrial remodeling. To test whether the impacts of HSF1 on mitochondria are unique to the aggressive PC3M cell line, we created another HSF1-KO model in the colon cancer cell line DLD1 (Fig. S2A). Loss of HSF1 in DLD1 cells also impaired cell proliferation but to a greater extent (increasing doubling time by ∼40%, Fig. S2B). Like the PC3M cells, the down-regulated proteins upon HSF1 knockout are most enriched among those in the mitochondrial matrix (Fig. S2C). In contrast to the PC3M cells, both mRNAs and proteins of mitochondrial genes disproportionately decreased upon HSF1 depletion (Fig. S2D&E), consistent with the larger defects in cell proliferation. Our data indicate that HSF1 is critical in regulating the steady-state mitochondrial proteome, which may underlie HSF1’s role in supporting the rapid proliferation of cancer cells.

### HSF1 promotes mitochondrial biogenesis and function

While the transcriptomic and proteomic data strongly suggest that, in the absence of HSF1, mitochondrial contents decrease in DLD1 cells, it remains unclear whether HSF1 is essential for overall mitochondrial biogenesis in PC3M cells. We performed kinetic analysis on mitochondrial protein synthesis using two complementary methods to address this question. First, we conducted stable isotope labeling using deuterium oxide (D2O) in the PC3M-WT and HSF1-KO cells over a 6-hour time course. Since deuterium can be incorporated into amino acids and nucleic acids, it labels both proteins and DNA (Fig. 3A)^36^. While the synthesis of DNA, cytosolic, and mitochondrial proteins is impaired in HSF1-KO cells (Fig. S3A-C), the decrease in mitochondrial protein synthesis is not merely due to slower cell proliferation, as the ratio of labeled mitochondrial proteins to DNA remains significantly lower in the HSF1-KO cells (Fig. 3B). In contrast, cytosolic protein synthesis aligns with the pace of DNA replication (Fig. 3C). These data indicate that mitochondrial biogenesis is impaired upon HSF1 depletion in PC3M cells. The results were confirmed by an experiment in which we labeled nascent proteins using the methionine analog, L-Homopropargylglycine (HPG), for a shorter duration (1 h). HPG specifically labels newly synthesized proteins, as confirmed by adding the translation inhibitor cycloheximide (CHX) (Fig. S3D). Western blot analysis indicates that we successfully enriched cytosolic and mitochondrial proteins in their corresponding fractions (Fig. 3D). While HPG labeling of cytosolic proteins is similar in PC3M-WT and HSF1-KO cells, newly synthesized proteins were reduced by about two-fold in the mitochondria in the absence of HSF1 (Fig. 3D). Consistent with the impaired mitochondrial biogenesis, respiration and ATP production also significantly decreased following HSF1 depletion from PC3M cells (Fig. 3E&F), suggesting that HSF1 may support cancer cell proliferation by enhancing mitochondrial biogenesis and energy metabolism.

**Fig. 3.**
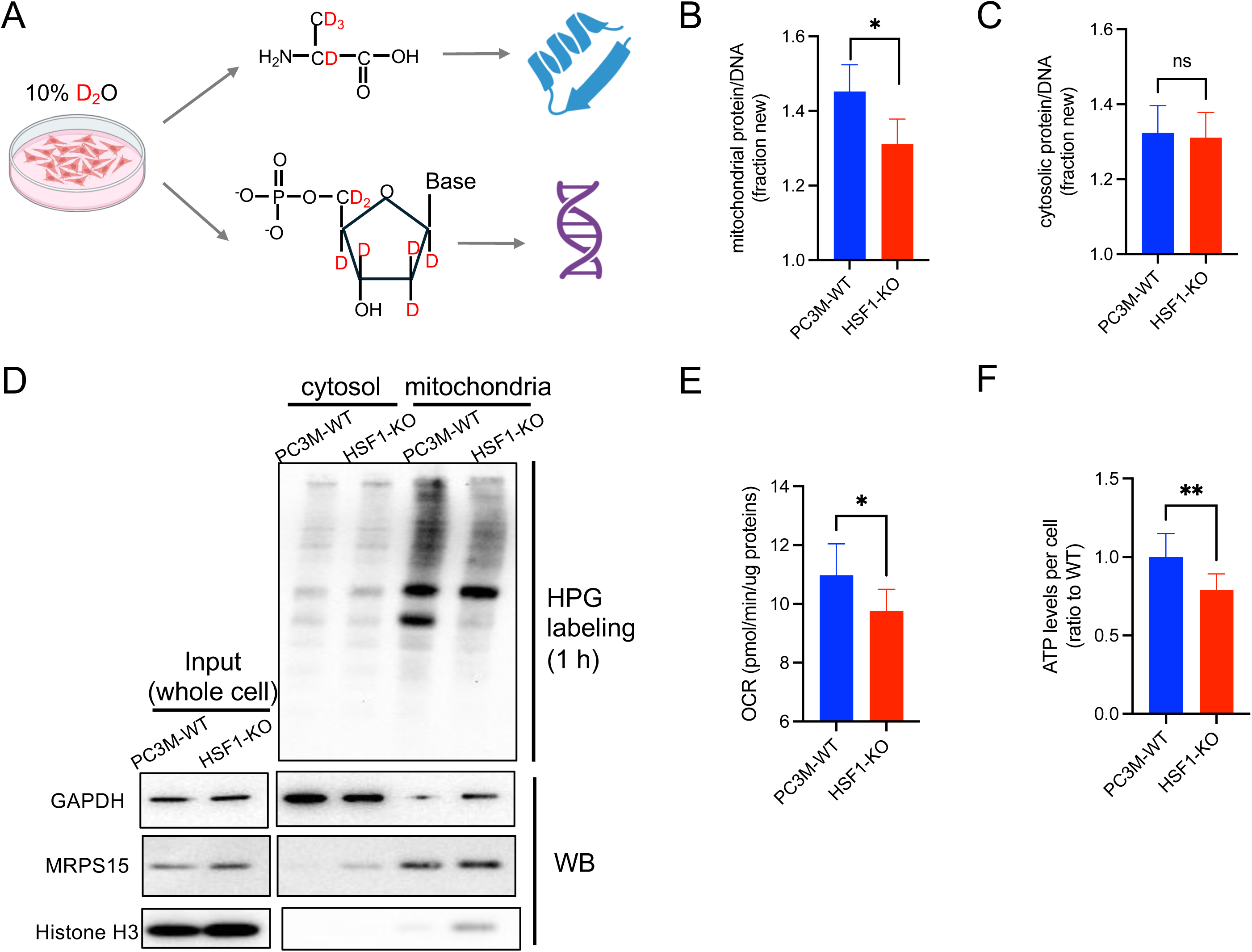
HSF1 promotes mitochondrial biogenesis and function. (A) Schematic diagram of deuterium oxide (D2O) labeling. (B&C) Histograms showing the ratios of newly synthesized mitochondrial proteins (B) and cytosolic proteins (C) to the newly synthesized DNA, as detected through deuterium oxide (D2O) labeling in PC3M-WT and HSF1-KO cells. Samples were collected hourly from 2 to 6 hours of labeling (n=3). The mean and SEM of the 5 time points are shown. Paired t-test: ns P>=0.05; * P<0.05. (D) Western blot analysis of newly synthesized cytosolic and mitochondrial proteins after one hour of HPG labeling in PC3M-WT and HSF1-KO cells. Upper panel: HPG-labeled proteins were biotinylated in a click reaction and detected using streptavidin-HRP and ECL. Lower panel: Western blot (WB) analysis of GAPDH, MPARS15, and Histone H3 as markers for cytosolic, mitochondrial, and nuclear proteins, respectively. (E&F) Basal oxygen consumption rate (OCR) (E) and cellular ATP levels (F) in PC3M-WT and HSF1-KO cells.

### HSF1 alters TIM23 core complex composition

Most mitochondrial proteins are encoded by nuclear genes, which are translated in the cytosol and imported into the mitochondria. Our data suggest that cytosolic translation rate is not reduced beyond the slowdown of cell proliferation (Fig. 3C&D), and mitochondrial gene mRNAs do not disproportionately decrease in PC3M cells upon loss of HSF1 (Fig. 2D). However, it is puzzling why more mitochondrial proteins are down-regulated than up-regulated at steady-state levels (Fig. 2C) and why mitochondrial biogenesis is significantly impaired (Fig. 3B&D). When comparing the transcriptomic and proteomic data, we found that 23 mitochondrial genes reduced protein levels at least 1.5-fold more than the changes in their mRNA levels upon HSF1 knockout (Fig. 4A). All of them are nuclear-encoded and reside in the matrix (18 out of 23) or the mitochondrial inner membrane (5 out of 23). The core complex of TIM23 translocase, comprising TIMM23 with TIMM17A or TIMM17B subunits, is essential for directing precursor proteins into the matrix or across the inner membrane ^37^. Proteomic data indicate that HSF1 knockout in PC3M cells leads to significant compositional changes in the TIM23 core complexes, in which TIMM17A decreases, TIMM17B increases, and TIMM23 remains unchanged (Fig. 4A). Western blot analyses confirmed this result (Fig. 4B&C). As TIMM17A is a confirmed genetic buffering interactor of HSF1, reduced TIMM17A may underlie the mitochondrial defects in HSF1-KO cells by impairing protein import through the TIM23 complex. Supporting this idea, TIMM17A KO is sufficient to reduce mitochondrial biogenesis (Fig. 4D) and entirely buffers the negative impacts of HSF1 KO on cell proliferation, respiration, and ATP production in PC3M cells (Fig. 4E-G). Thus, HSF1 enhances mitochondrial biogenesis and cell proliferation in a TIMM17A-dependent manner.

**Fig. 4.**
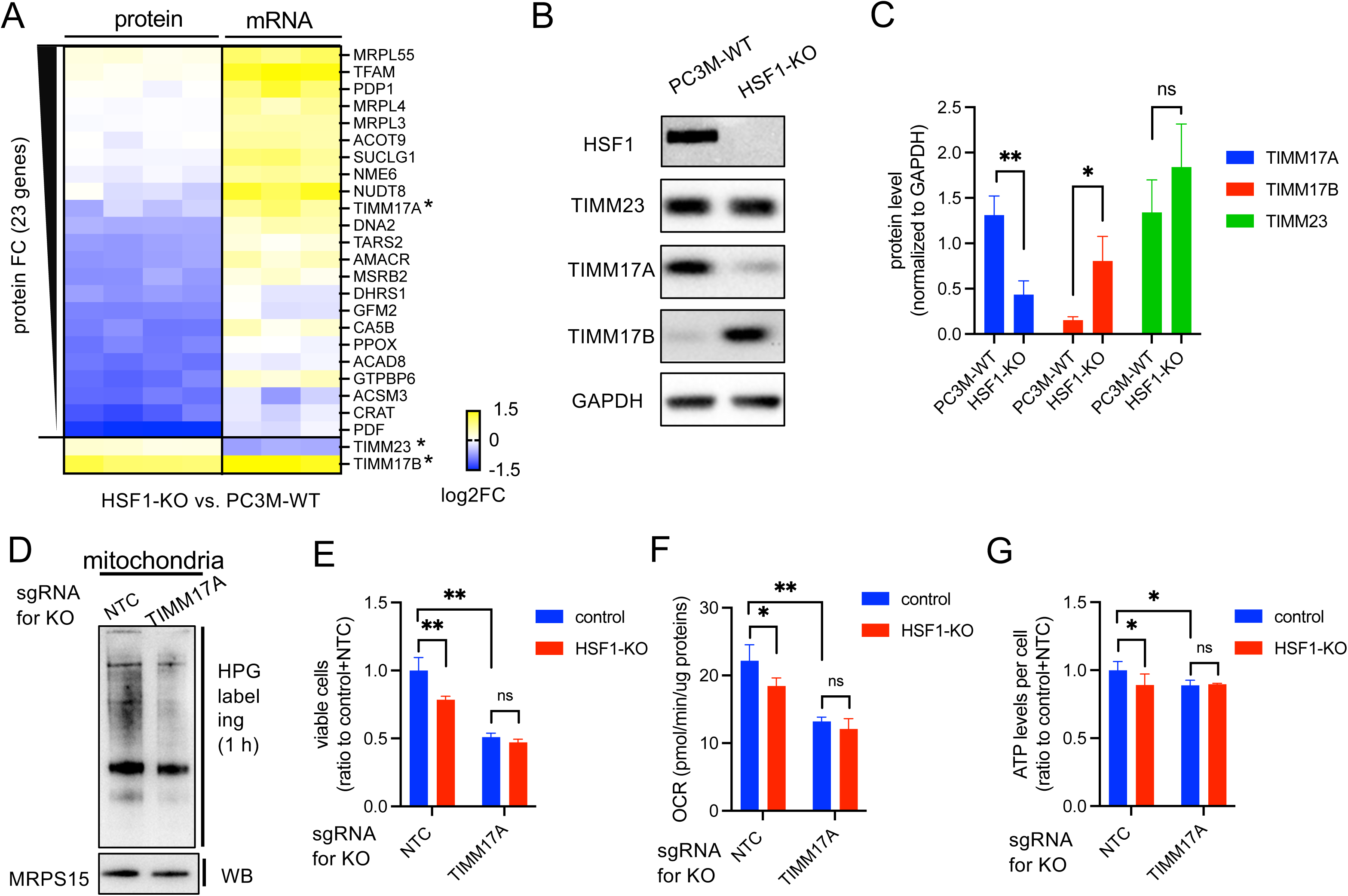
HSF1 alters TIM complex composition. (A) Heatmap showing log2 fold change (FC) of protein and mRNA of selected mitochondrial genes upon knockout of HSF1 in PC3M cells. The 23 mitochondrial genes that reduced protein levels at least 1.5-fold more than the changes in their mRNA levels are shown, along with the two additional subunits of the TIM23 core complex. The asterisks indicate the subunits of the TIM23 core complex. (B&C) Representative images (B) and the quantification (mean ± SD, n=3) (C) of western blot analysis on the TIM23 core complex in PC3M-WT and HSF1-KO cells. (D) Newly synthesized mitochondrial proteins after one hour of HPG labeling in the PC3M cells with TIMM17A knockout or treated with a non-target control (NTC). Western blot (WB) analysis of MRPS15 serves as a loading control for steady-state mitochondrial proteins. (E-G) Histograms displaying the number of viable cells (E), basal oxygen consumption rate (OCR) (F), and cellular ATP levels (G) in PC3M cells with single or double knockout of HSF1 and TIMM17A. After antibiotic selection, cells were seeded on Day 0. Viable cells were measured by PrestoBlue assays on Day 4. OCR and ATP were measured on Day 1.

### HSF1 regulates TIMM17A protein via the mitochondrial chaperonin complex

We then explore how TIMM17A protein levels are linked to HSF1 activity. We determined the HSF1 transcriptional program by combining CUT&Tag to map the HSF1 genomic binding sites and RNA-seq to measure changes in mRNA in PC3M-WT and HSF1-KO cells. In agreement with previous reports that HSF1 directly controls a compact transcriptional program^7,20^, we identified 173 HSF1 binding peaks in the PC3M-WT cells (Fig. 5A), which are enriched with the HSF1 binding motif (Fig. 5B). Experiments in the HSF1-KO cells produced very few sequencing reads, verifying the specificity of CUT&Tag. Among the 97 protein-coding genes associated with HSF1 peaks within 2 kb of the transcription start sites, 37 genes significantly altered their mRNA levels following HSF1 knockout (Fig. 5C), in which 34 are down-regulated, consistent with the established role of HSF1 as a transcriptional activator. The head-to-head HSPD1 and HSPE1 genes are among the HSF1 direct targets that show the most significant reduction in mRNA levels upon HSF1 depletion (Fig. 5C&D), leading to a ∼60% decrease in protein levels (Fig. 5E). HSPD1 and HSPE1 encode the HSP60 and HSP10 chaperones that form the mitochondrial chaperonin complex, responsible for refolding proteins once imported into the matrix ^38^. We wonder if TIMM17A protein levels are sensitive to the protein folding capacity in the matrix. We knocked down HSPD1/HSP60 using siRNA and monitored the mitochondrial chaperonin and TIMM17A over time (Fig. 5F-H). In our experimental condition, HSPD1 RNAi did not significantly impair proliferation until 96 h (Fig. 5F). HSPD1 protein was reduced by 50% at 24 h of siRNA treatment and reached a similar level as in the HSF1-KO cells at 48 h. HSPE1 protein also started declining at 24 h but remained at about the 75% level. This could be due to induced HSPE1 expression by the mitochondrial unfolded protein response (mtUPR) in the presence of HSF1 and may be related to the HSPD1-independent role of HSPE1 ^39^. TIMM17A protein displayed a delayed decline upon HSPD1 knockdown and significantly decreased at 72 h, supporting the idea that reduced TIMM17A results from the compromised function of the mitochondrial chaperonin complex.

**Fig. 5.**
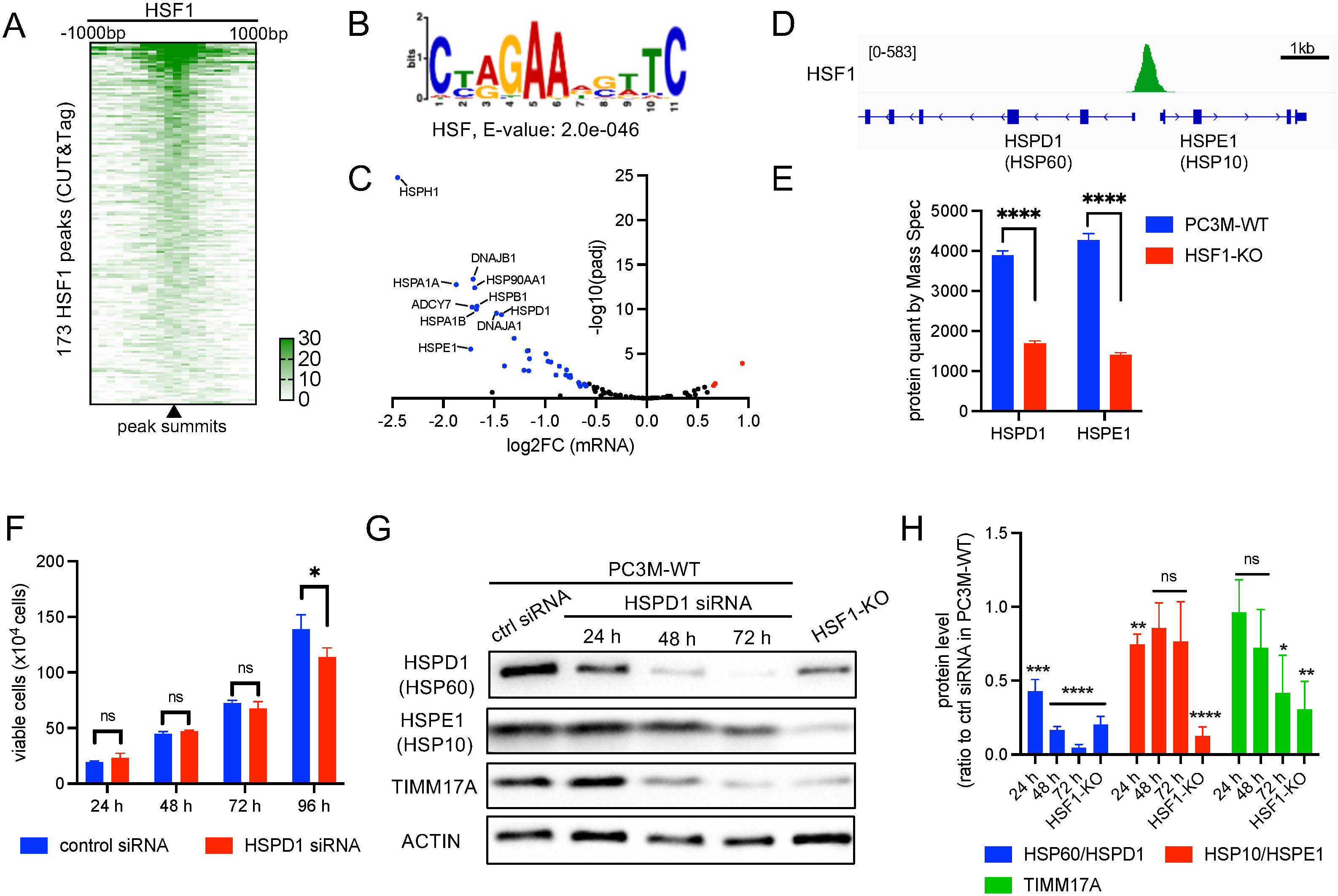
HSF1 regulates TIMM17A protein via the mitochondrial chaperonin complex. (A) Heatmap of HSF-1 occupancy at CUT&Tag peaks in PC3M cells. Normalized HSF-1 reads were mapped to 100 bp bins, ±1000 bp from the peak summits. (B) The top DNA motif enriched at the center of HSF1 CUT&Tag peaks (±100 bp from the peak summits). (C) Scatter plots of mRNA log2 fold change (FC) at HSF1-bound genes upon HSF1 knockout in PC3M cells. The 97 genes associated with HSF1 CUT&Tag peaks within 2kb of the transcription start sites are shown. Differentially expressed (DE) genes (1.5-fold, FDR: 0.05) are indicated in blue (down-regulated) or red (up-regulated) color. The top 10 DE genes with the most significant fold change are labeled with gene names. (D) Genome browser view of HSF1 occupancy at the head-to-head HSPD1 and HSPE1 genes. (E) Protein levels of HSPD1 and HSPE1 determined by TMT proteomics in PC3M wild-type cells (PC3M-WT) and PC3M cells with HSF1 knockout (HSF1-KO). Unpaired t-test: **** P < 0.0001. (F) Histograms showing the number of viable cells upon HSPD1 siRNA treatment in PC3M cells. (G&H) Representative images (G) and quantification (mean ± SD, n=3) (C) of western blot analysis on HSPD1, HSPE1, and TIMM17A upon HSPD1 siRNA treatment. In each experiment, the tested proteins were first normalized to the ACTIN loading control, and the ratios relative to the protein level in the control siRNA (ctrl) were calculated and plotted into histograms. Unpaired t-test (against control siRNA): ns P >= 0.05; * P < 0.05; ** P < 0.01; *** P < 0.001; **** P < 0.0001.

### TIMM17A promotes mitochondrial translation

Multiple mitochondrial matrix proteins were decreased at protein levels more significantly than mRNA levels upon loss of HSF1 in PC3M cells (Fig. 4A). Among them are subunits of mitochondrial ribosomes (MRPL55, MRPL4, and MRPL3) and essential regulators of mitochondrial translation, including TARS2 (the mitochondrial threonyl-tRNA synthetase), GFM2 (a mitochondrial ribosome recycling factor), and PDF (mitochondrial peptide deformylase). We wonder whether HSF1 supports mitochondrial biogenesis partially by maintaining the levels of the mitochondrial translation machinery. Using commercially available antibodies, we confirmed with western blot analysis that TARS2 and GFM2 proteins decreased upon HSF1 knockout in PC3M cells (Fig. 6A&B). These genes did not significantly decrease their mRNA levels upon HSF1 depletion (Fig. 4A), indicating that regulation occurs at the post-transcriptional level. The TARS2 antibody recognized a higher molecular band that increased upon HSF1 knockout (Fig. 6A). Although it is tempting to think that this band may represent the precursor protein of TARS2, the TARS2 knockout experiment suggests otherwise (Fig. 6C). Co-essentiality analysis suggests that TIMM17A has a much stronger positive correlation with the mitochondrial translation machinery than TIMM17B does (Fig. 6D&E, Kolmogorov-Smirnov test for TIMM17A vs. TIMM17B: P=9.92e- 43). We then wonder if HSF1’s impacts on TARS2 and GFM2 may be through TIMM17A. Knockout of TIMM17A led to a significant decrease in TARS2 protein but not GFM2 (Fig. 6F&G). Instead, both TARS2 and GFM2 proteins were reduced upon HSPD1 siRNA (Fig. S5A&B). Additionally, TIMM17A KO did not change HSPD1 and HSPE1 protein levels, consistent with TIMM17A functioning downstream of the mitochondrial chaperonin complex in HSF1-mediated mitochondrial remodeling (Fig. 5G&H). In contrast, the TIMM17B protein increased upon TIMM17A knockout. Collectively, our data suggest that HSF1 regulates TARS2 and TIMM17B at least partially through TIMM17A, while it regulates GFM2 in a HSPD1/HSPE1-dependent but TIMM17A-independent manner.

**Fig. 6.**
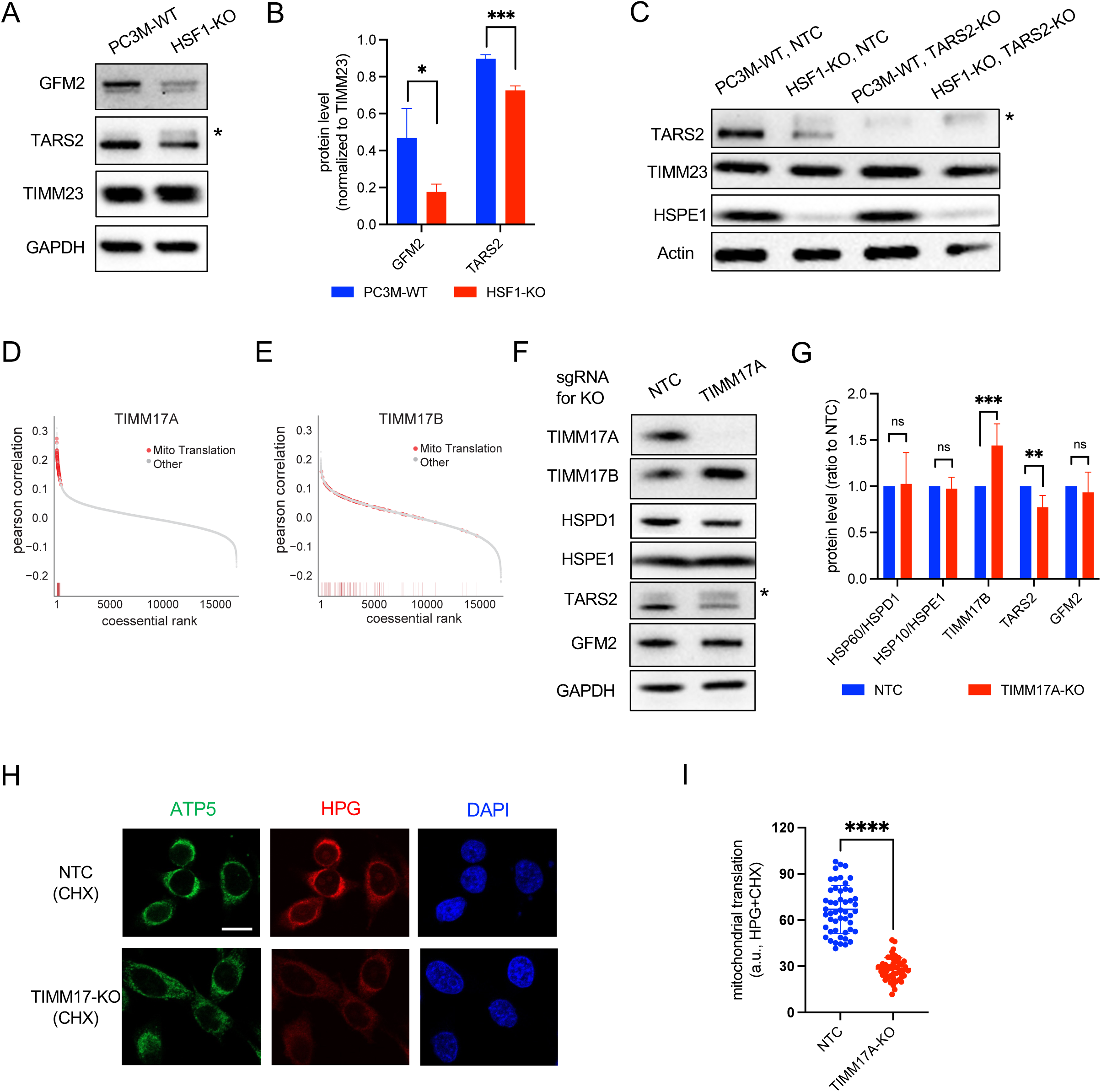
TIMM17A promotes mitochondrial translation. (A&B) Representative images (A) and quantification (mean ± SD, n=3) (B) of western blot analysis for mitochondrial translation factors, GFM2 and TARS2, in PC3M wild-type cells (PC3M-WT) and PC3M cells with HSF1 knockout (HSF1-KO). The asterisk indicates a higher molecular band recognized by the TARS2 antibody. Unpaired t-test: * P < 0.05; *** P < 0.001. (D) Western blot analysis of TARS2 knockout (TARS2-KO) in PC3M wild-type cells (PC3M-WT) and PC3M cells with HSF1 knockout (HSF1-KO). NTC: non-target control. The asterisk indicates a likely non-specific band recognized by the TARS2 antibody. (D&E) Coessentiality rank and correlation coefficient for all genes with TIMM17A (D) and TIMM17B (E). The positions of mitochondrial translation factors (Mito Translation) in the coessentiality plots are indicated as red dots. (F&G) Representative images (F) and quantification (mean ± SD, n=3) (G) of western blot analysis in PC3M cells following TIMM17A knockout. The asterisk denotes a likely non-specific band recognized by the TARS2 antibody. Either MAPT1 (n=4) or AAVS (n=2) was employed as the negative control in the co-CRISPR assays along with a non-target control (NTC) sgRNA. In each experiment, the tested proteins were normalized to the GAPDH loading control, and the ratios relative to the protein level in NTC were calculated. Results (n=6) were pooled for quantification and plotted into histograms. Unpaired t-test: ns P >= 0.05; ** P < 0.01; *** P < 0.001. (H&I) Representative images (H) and quantification (mean ± SD) (I) of HPG labeling for newly synthesized proteins by mitochondrial translation. HPG labeling was performed in PC3M cells with TIMM17 knockout (TIMM17-KO) or the non-target control (NTC) for 45 min in L-methionine- free medium containing cycloheximide (CHX) to inhibit cytosolic translation. MAPT1 was used as the negative control in the co-CRISPR assay. Immunofluorescence of ATP5 was used as a mitochondrial marker, and DAPI was used to stain DNA. Scare bar: 20 μM. Unpaired t-test: **** P < 0.0001.

Finally, we tested the impact of TIMM17A on mitochondrial translation through HPG labeling in the presence of cycloheximide as previously described ^40^. We confirmed that cycloheximide inhibited translation in the cytosol while allowing mitochondrial translation (Fig. S5C&D). Using either MAPT1 or AAVS as the negative control for CRISPR, and employing two different mitochondrial markers, our results indicate that TIMM17A knockout significantly reduced mitochondrial translation (Fig. 6H&I and S5E&F). Our findings suggest that TIMM17A could sense protein folding capacity in the matrix and adjust mitochondrial translation.

### TIMM17A protein level is coupled with HSF1 activity to promote robust cell proliferation

TIMM17A has a short half-life and is sensitive to proteotoxic stress that attenuates cytosolic translation ^41^. We have confirmed the short half-life of TIMM17A in PC3M cells with cycloheximide treatment followed by TMT-proteomics and western blot analyses (Fig. S6A&B). Interestingly, 6 hours of cycloheximide treatment also significantly decreased TIMM23 protein levels without changing TIMM17B (Fig. S6A). This is consistent with the recent report that the TIMM23 and TIMM17A-containing translocase channel is degraded if unoccupied by precursor proteins, while TIMM17B is stable ^42^. Under stress and physiological conditions, the ATP-dependent protease YME1L1 is responsible for TIMM17A degradation ^41–44^. We then ask whether increasing TIMM17A protein levels by inhibiting YME1L1-mediated degradation in HSF1-KO cells will restore cell proliferation or be more detrimental. Knockdown of YME1L1 by siRNA increased TIMM17A protein levels by about 2-fold in HSF1-KO cells (Fig. 7A) and exhibited clear synthetic lethality with HSF1 knockout (Fig. 7B&C). This result suggests that degradation of TIMM17A by YME1L1 serves as a protective response, tuning down protein import when folding capacity is limited in the matrix.

**Fig. 7.**
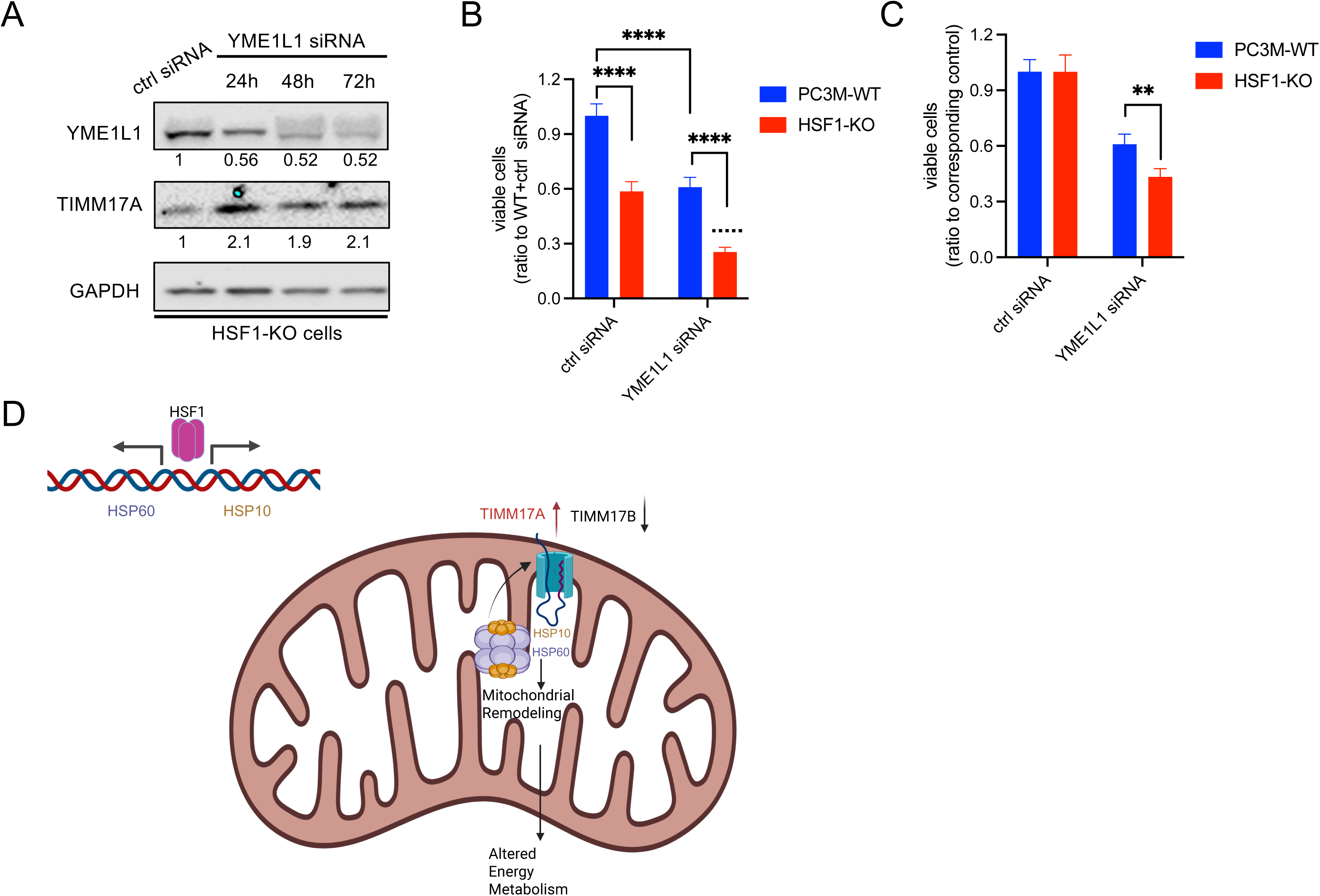
TIMM17A protein level is coupled with HSF1 activity to promote robust cell proliferation. (A) Western blot analysis of TIMM17A upon YME1L1 knockdown. PC3M cells with HSF1 knockout (HSF1-KO) were treated with YME1L1 siRNA or control (ctrl) siRNA. YME1L1 and TIMM17A protein levels were monitored over time with GAPDH as a loading control. The tested proteins were normalized to GAPDH, and the ratios relative to those in the control siRNA are labeled. (B&C) Histograms showing the numbers of viable cells measured by PrestoBlue assays on Day 5 after siRNA treatment. The wildtype PC3M cells (PC3M-WT) and HSF1-KO cells were treated with YME1L1 siRNA or control (ctrl) siRNA. The ratios to the cell number in PC3M-WT + control siRNA (B) and the ratios to the cell numbers in the corresponding control siRNA (either PC3M- WT or HSF1-KO) (C) are presented (mean ± SD, n=3). Dashed lines represent the calculated additive effects. Unpaired t-test: ** P < 0.01; **** P < 0.0001. (E) A proposed model for HSF1-mediated mitochondrial remodeling via regulating the TIM23 core complex composition.

## Discussion

In this study, we determined the epistatic interaction network of HSF1 in cancer cells through paired CRISPR screens in the wildtype PC3M and HSF1 knockout cells, followed by individual validation using our co-CRISPR assay. These HSF1 interactors are enriched with subunits of protein complexes (e.g., Mediator, CPSF, GINS, RFC, and CHAF1) and proteins functioning in the same pathways (Fig. 1H), demonstrating the specificity of our functional genomics analysis. Our results support the reported roles of HSF1 in protein folding ^10,11^, DNA repair ^12^, and protein synthesis ^13,14^, and have revealed previously underappreciated cellular functions that HSF1 may contribute to. First, HSF1 interactors are highly enriched with regulators of transcription elongation. Most of them show genetic buffering effects on HSF1, including the Mediator complex, which activates transcription initiation and elongation ^45^, and multiple transcription elongation factors that regulate Pol II pausing (NELFCD and SUPT4H1), pause release (SUPT4H1 and ELL), and passage through chromatin (SSRP1) ^46^. In addition, HSF1 exhibits a synthetic lethal interaction with HEXIM1 that inhibits the P-TEFb kinase responsible for productive elongation ^47^. It is possible that these factors co-regulate HSF1 target genes. However, given that they control transcription elongation globally at many genes, the genetic interaction is unlikely to be dictated by the small transcriptional program that HSF1 directly regulates. Future studies will determine if HSF1 has a role that impacts global transcription elongation in cancer cells. Furthermore, HSF1 genetically interacts with genes that influence multiple aspects of mitochondrial activities (Fig. 1H). Our data suggest that HSF1 significantly remodels the mitochondrial proteome by altering TIM23 translocase composition, and regulates mitochondrial biogenesis, ATP production, and cell proliferation in a manner dependent on TIMM17A, a subunit of the core TIM23 complex.

Despite the Warburg effect, mitochondria play an essential role in cancer cells. A recent study shows that disrupting mitochondrial proteostasis effectively inhibits cell growth and tumor progression in prostate cancer ^48^. HSF1 activates mtUPR, a cellular stress response that safeguards mitochondrial proteostasis ^49^, and regulates glucose and NAD(+) metabolism, impacting mitochondrial activities in cancer cells ^15,16^. However, the role of HSF1 in mitochondrial biogenesis in cancer cells and the underlying mechanisms were not understood. Our data suggest HSF1 promotes mitochondrial biogenesis by regulating the TIM23 core complex composition (Fig. 4 and S4). The protein level of TIMM17A, a short-lived subunit of the TIM23 complex ^41^, depends on HSPD1 and HSPE1, the direct target genes of HSF1 that encode the mitochondrial chaperonin complex. When HSF1 activity is high, the ratio of TIMM17A to its paralog TIMM17B is high; while in HSF1-KO cells, TIMM17A protein levels decrease, likely due to degradation by YME1L1. Even though TIMM17B increases upon loss of HSF1 or TIMM17A and the TIMM23 protein level maintains in PC3M cells, it is insufficient to restore mitochondrial biogenesis, respiration, and ATP production (Fig. 7D). These results suggest an essential role for TIMM17A in cancer cells, consistent with a recent report that TIMM17A is overexpressed in lung cancer and associated with poor prognosis ^50^. As TIMM17A is short-lived (Fig. S6A), TIMM17B has been considered the housekeeping TIM17 subunit. However, our data in TIMM17-KO cells suggest that the feedback up-regulation of TIMM17B is insufficient to restore mitochondrial protein uptake (Fig. 4D). While this could be due to a difference in expression levels, it is more tempting to think that TIMM17A and TIMM17B may have a preference in precursor proteins that they import. We found that in HSF1-KO cells, mitochondrial translation factors selectively decrease at the protein levels (Fig. 4A). Published proteomics analyses have shown that the mitochondrial translation machinery is among those most sensitive to import efficiency ^51^. At least one of these translation factors and a genetic interactor of HSF1, TARS2, alters its protein level in a TIMM17A-dependent manner (Fig. 6F&G). A recent structural study suggests that TIM17 instead of TIM23 forms the translocase channel ^52^, likely providing the import specificity. Despite TIMM17A and TIMM17B having very similar predicted structures ^53^, biochemical studies showed that TIMM17A and TIMM17B- containing complexes exhibit different import rates based on candidate precursor proteins ^54^. It is interesting for future studies to determine if TIMM17A and TIMM17B contribute to mitochondrial import specificity, especially toward mitochondrial translation factors.

Regulation of mitochondrial import is essential for cellular proteostasis, as it not only controls mitochondrial biogenesis but also precursor accumulation in the cytosol, which leads to proteotoxic stress ^55,56^. TIMM17A has been shown to sense cytosolic stress that attenuates cytosolic translation and contributes to mtUPR ^41^. Our data suggest that TIMM17A could also serve as a sensor for folding capacity in the mitochondrial matrix and adjust mitochondrial biogenesis accordingly. We show that depletion of TIMM17A by YME1L1 is a protective response when HSF1-dependent HSPD1/HSPE1 expression is low (Fig. 7B&C). We did not observe significant changes in other YME1L1 substrates as identified in the hypoxia response ^43^ (Table S2), suggesting degradation of TIMM17A in HSF1-KO cells is different from that in stress response, consistent with a very recent report that YME1L1 could also degrade TIMM17A in physiological conditions ^42^. HSF1 is reported to regulate YME1L1 expression in adhesion-mediated mechanosignaling ^57^. However, we did not observe YME1L1 changes at protein or mRNA levels in HSF1 knockout cancer cells (Tables S2&S3). Additional mitochondrial proteins could be involved in the folding capacity sensing by TIMM17A and YME1L1. These include the PAM complex, which serves as the motor for TIM23-mediated protein import and consists of mitochondrial HSP70 and HSP40 chaperones ^37^, as well as OCIAD1 and prohibitins that are shown to regulate the stability of TIMM17A in a recent publication ^44^. Future studies will dissect the interplays among these factors, mitochondrial chaperonin, and YME1L1 to reveal the underlying mechanisms.

## Methods

### CRISPR screen and data analysis

Virus production and infection were performed as previously described ^58^. Briefly, lentivirus was obtained by co-transfecting the Brunello libraries, packaging vector psPAX2, and the envelope vector pVSV-G (Addgene) into 293FT cells in FBS-depleted media using Lipofectamine 3000 (ThermoFisher) following the manufacturer’s instructions. After 24 hours, the media was replaced with fresh media containing 10% FBS and 1% Pen-Strep. After 72 hours of transfection, 25 mM HEPES (pH 7.4) was added, and lentivirus was harvested by centrifugation at 500 x g for 10 minutes. Supernatants were collected, and a small-scale infection was performed to determine the MOI following the published procedure ^58^. The virus was titrated to MOI (0.5-1), and infection was performed in 25 *10-cm plates. This gave ∼250-fold coverage of the libraries. Cells were selected by 2ug/ml puromycin for three days to establish the initial population (Day 0). Cells were then split and collected every 3 to 4 days. The first pair of screens was performed for 15 days, and the second pair lasted 20 days to increase the sensitivity. Genomic DNA was isolated using QIAamp DNA Blood Maxi Kit according to the manufacturer’s instructions. sgRNAs were amplified by PCR following the Broad Institute protocol using ExTaq polymerase, and the PCR products were purified using AMPure XP beads before sending for Illumina sequencing.

The sgRNAs were quantified by building bowtie indices from the Brunello libraries’ sgRNA information spreadsheet and mapping the sequencing reads to the indices using bowtie2’s local mode. Only the reads that perfectly matched the forward strand were kept. The sgRNA quantification across the time course of negative selection was fit to the exponential decay model using homemade scripts to calculate the beta value (CRISPR score). Statistical tests were performed against the non-target controls to determine the essential genes (Z-score cutoff: 4), and between PC3M-WT and HSF1-KO to determine genetic buffering and synthetic lethal interactors. See details in the supplemental method.

### RNA-seq analysis

RNA was isolated and purified using the Quick-RNA MiniPrep Kit (Zymo Research), following the manufacturer’s instructions, using on-column DNase I digestion to remove genomic DNA. Total RNAs were polyA enriched, and directional RNA-seq libraries were prepared using NEBNext Ultra II RNA library prep Kit. Sequencing was done at a NovaSeq 6000 sequencer at the OMRF clinical genomics core. RNA-seq reads were mapped to the GRCh38 (hg38) genome using RNA STAR^59^. The mapped reads were then subject to FeatureCounts in Rsubread ^60^ for quantification. Differential expression (DE) analyses were then done using DESeq2 ^61^ with default settings.

### CUT&Tag analysis

CUT&Tag was performed in PC3M-WT and HSF1-KO cells in duplicates using the CUTANA™ CUT&Tag Kit following the manufacturer’s instructions, except that a secondary antibody was added to increase the reads’ sequence diversity to avoid issues during sequencing reactions. HSF1 antibody (Rabbit, Cell Signaling) was used. The sequencing libraries were prepared by PCR from the genomic DNA using the primers against the tagged sequencing adaptors. Pair-end sequencing was performed on the MiSeq sequencer at the OMRF clinical genomics core.

Sequencing reads were mapped to the GRCh38 (hg38) genome using bowtie2 ^62^. Duplicate reads were filtered for each replicate using MACS2 ^63^. HSF1 peaks were called using MACS2 for each replicate, and the shared peaks in the biological duplicates were kept for subsequent analyses. To visualize the CUT&Tag data in genome browser views, the bedgraph files were normalized to reads per million using MACS2 callpeak -B –SPMR, and visualized using Integrative Genomics Viewer (IGV) ^64^. The motif analysis was done using MEME-ChIP ^65^ on the regions 100 bp around the peak summits.

### Click-Chemistry Labeling of Mitochondrial Translation

To monitor mitochondrial translation, we performed Click-Chemistry Labeling of Mitochondrial Translation, adapted from Zorkau et al., with minor modifications ^66^. Click-iT® HPG Alexa Fluor® Protein Synthesis Assay Kit was used for detection. Cells grown on glass-bottom dishes were labelled with 50 µM HPG in pre-warmed L-methionine-free medium for 45 minutes in the presence of 50 μg/mL cycloheximide to inhibit cytosolic translation. To eliminate unincorporated HPG, cells were pre-permeabilized for 80 seconds at room temperature using 0.005% digitonin in mitochondria-protective buffer (MPB; 10 mM HEPES/KOH, 10 mM NaCl, 5 mM MgCl2, and 300 mM sucrose in H2O, pH 7.4). Subsequently, cells were washed once with MPB and fixed in pre-warmed MPB containing 8% paraformaldehyde for 7 minutes. After washing twice with PBS with 3% BSA, cells were fully permeabilized using 0.5% (v/v) Triton X-100 in 1X PBS (pH 7.4). Cells were then blocked with PBS with 5% BSA for 10 minutes, followed by a washing step using PBS with 3% BSA before a 30-minute incubation with the Click-iT reaction cocktail per the manufacturer’s instructions. Immunofluorescence was performed to locate mitochondria. Following one washing step with the Click-iT rinse buffer, MRPS15 or ATP5 primary antibody (Proteintech) was added (1 to 400 dilution) in PBS with 5% BSA and incubated for 1 hour. Goat anti-rabbit IgG secondary antibody conjugated with Alexa Fluor Plus 488 (ThermoFisher) was added to detect MRPS15 or ATP5. DNA staining was performed in 250 ng/mL DAPI for 15 minutes at room temperature. Samples were visualized using a Zeiss LSM980 Confocal Microscope with a 20X objective.

### Nascent protein labeling and mitochondria import assay

Mitochondrial fractionation was performed following a previous report ^67^ with minor modifications. In brief, methionine was depleted by incubating cells in pre-warmed L-methionine-free medium for 20 minutes before 1-hour incubation of 50 µM HPG in the presence of 50 μg/mL cycloheximide. Cells were collected by trypsinization and homogenized with a small-clearance pestle B in isotonic buffer (250 mM sucrose, 1 mM EDTA, 10 mM Tris-HCl, pH 7.4). Mitochondria were separated from cytosol by differential centrifugation. Homogenate was centrifuged at 600 x *g* for 15 min, and the supernatant was spun at 10,000 x *g* for 25 minutes to separate the mitochondrial pellet and the cytosolic supernatant. To purify mitochondria, the mitochondrial pellet was washed once in an isotonic buffer (250 mM sucrose, 1 mM EDTA, 10 mM Tris-HCl, pH 7.4), followed by centrifugation at 500 x *g* for 5 minutes. The supernatant was centrifuged at 10,000 x *g* for 25 minutes to attain a purified mitochondrial pellet, which was resuspended in 0.4 % SDS containing PBS. The cytosolic supernatant was further purified by centrifugation at 18,000 x *g* for 30 minutes, and the supernatant was collected. The cytosol fraction was solubilized with 0.4% SDS in PBS. Protein concentration was determined by BCA assays (ThermoFisher). Approximately 40 µg protein samples were solubilized in 60 µL of 0.4 % SDS in PBS for 10 minutes at room temperature, followed by centrifugation at 18,000 x *g* for 5 minutes. Supernatants were incubated with click reaction cocktail (20 µM Biotin Picolyl Azide, 1.2 mM BTTAA, 600 µM CuSO4, and 5 mM (+)- Sodium L-ascorbate) at a final concentration of 0.2% SDS for 1 hour. Proteins were precipitated by the chloroform-methanol extraction method ^68^, and the obtained pellets were resolubilized in 1X SDS loading buffer (Sigma).

### siRNA transfection

Gene silencing by small interfering RNA (siRNA) was performed using Lipofectamine RNAiMAX (Invitrogen). Diluted siRNA and diluted Lipofectamine® RNAiMAX Reagent were prepared separately and incubated for 5 minutes at room temperature. Specifically, 2.5 µL of 10 µM ON-TARGETplus Non-targeting Control Pool or Human HSPD1 siRNA (Dharmacon) was added to 250 µL of optiMEM media. On the other hand, 5 µL of Lipofectamine® RNAiMAX Reagent was diluted in 250 µL of optiMEM media. The diluted siRNA and Lipofectamine® RNAiMAX Reagent were mixed and incubated for 15 minutes at room temperature before dropwise addition into cells at a final concentration of 10 nM siRNA. Cells were collected 24, 48, and 72 hours after transfection by trypsinization and lysed in RIPA buffer (Sigma-Aldrich) containing Protease inhibitor cocktail (Roche) for western blot analysis.

### Deuterium oxide (D2O) labeling

The experiments were performed as previously described ^69^ with modifications. Briefly, labeling was performed in media with 10% D2O over a 6-hour course. Cells were collected hourly after 2 hours of labeling. Cytosolic, mitochondrial, and nuclear fractions were isolated from cells using differential centrifugation to prepare cytosolic protein, mitochondrial protein, and DNA. The incorporation of deuterium into alanine and deoxyribose was analyzed by GC-MS to calculate the fractional synthesis of protein and DNA. See details of experimental procedure and data analysis in the supplemental information.

### Western blot

Cells were lysed with RIPA buffer (Sigma-Aldrich) containing protease inhibitor cocktail (Roche). Proteins were separated by 4 -12% NuPAGE Bis-Tris protein gels (ThermoFisher) and transferred onto 0.45 µm nitrocellulose blotting membrane. Membranes were blocked with 3% nonfat milk for 3 hours and incubated with primary antibodies overnight in 3% nonfat milk, followed by a 2-hour incubation with secondary antibodies. Proteins were visualized using Clarity Western ECL Substrate (Bio-Rad) and imaged by iBright CL1500 Imaging System (ThermoFisher). Signals were quantified using the iBright Analysis Software.

### PrestoBlue assay

Cell proliferation and viability were measured using PrestoBlue™ Cell Viability Reagents according to the manufacturer’s instructions. In brief, cells were incubated with the Prestoblue reagent (ThermoFisher) for 30 minutes. Fluorescent signals were detected by FLUOstar Omega (BMG Labtech) with excitation and emission wavelengths at 544 and 590 nm, respectively.

## Data and Code Availability

The sequencing data, proteomics data, and the code for CRISPR screen analysis will be uploaded to GEO, PRIDE, and GitHub, respectively, and made public upon publication.

## Acknowledgement

We thank Xiaowen Wang for sharing the mitochondrial fractionation protocol. We thank the Imaging Core Facility at NYMC for assistance with confocal imaging, the OMRF clinical genomics core for RNA-seq and Cut&Tag sequencing, and the ThermoFisher Center of Multiplexed Proteomics at Harvard for total protein quantification. This work is supported by the NIH grant R35GM138364 to JL.

