## supplemental figures and legend for "HSF1 remodels mitochondrial biogenesis and function in cancer cells via TIMM17A"

Fig.S1_Nguyen et al.

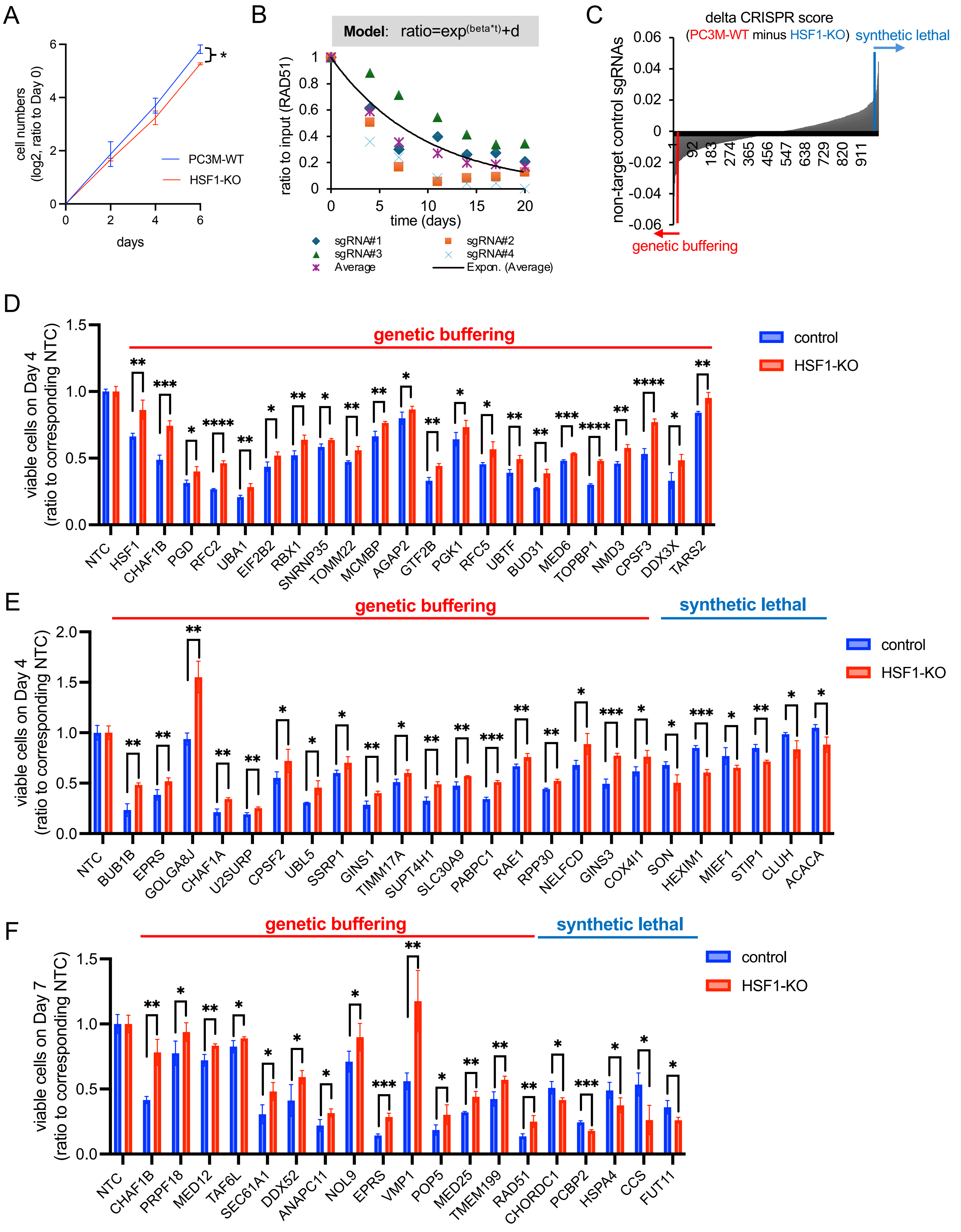

G

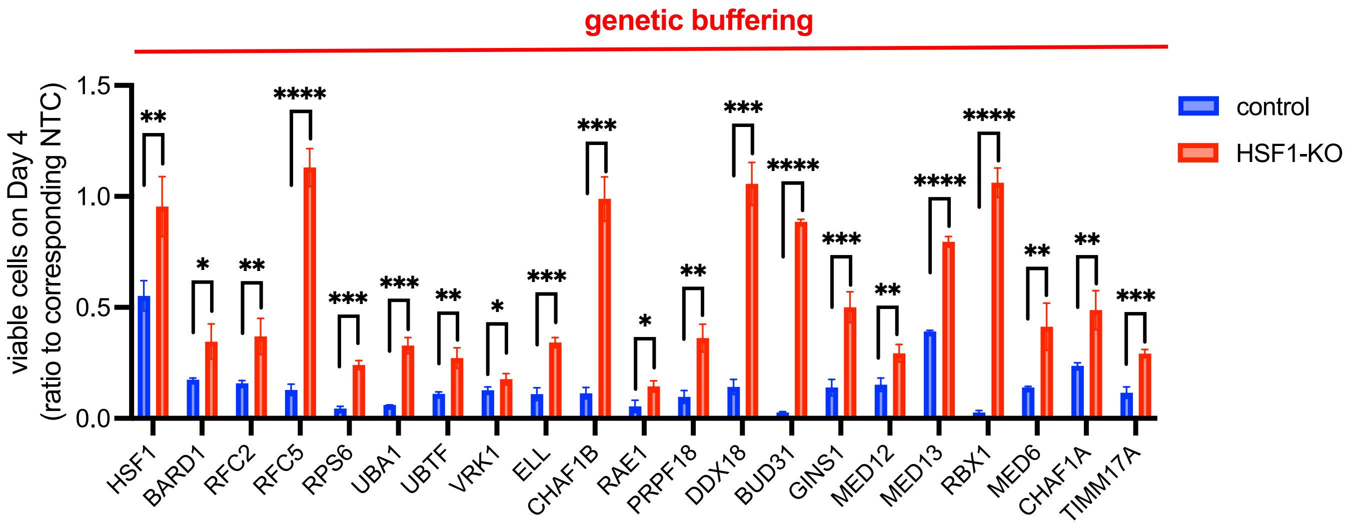
genetic buffering
viable cells on Day 4
(ratio to corresponding NTC)
1.5
1.0
0.5
0.0
control
HSF1-KO
HSF1
BARD1
RFC2
RFC5
RPS6
UBA1
UBTF
VRK1
ELL
CHAF1B
RAE1
PRPF18
DDX18
BUD31
GINS1
MED12
MED13
RBX1
MED6
CHAF1A
TIMM17A

H

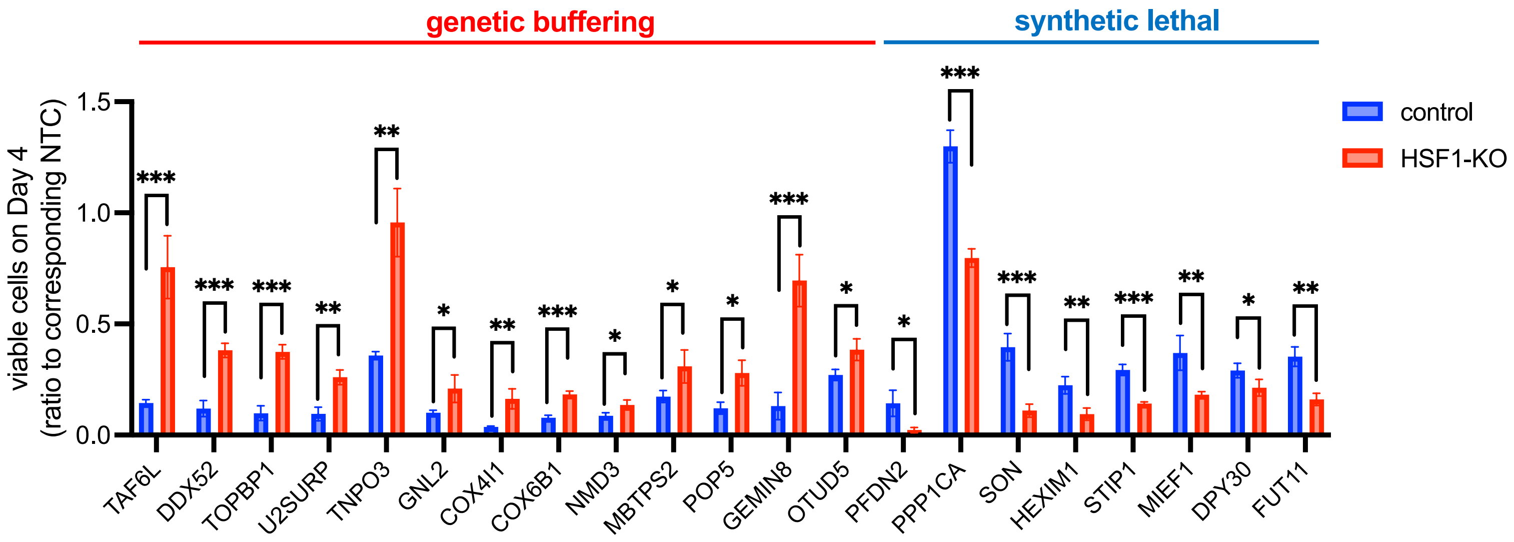
genetic buffering
synthetic lethal
viable cells on Day 4
(ratio to corresponding NTC)
1.5
1.0
0.5
0.0
control
HSF1-KO
TAF6L
DDX52
TOPBP1
U2SURP
TNPO3
GNL2
COX4I1
COX6B1
NMD3
MBTPS2
POP5
GEMIN8
OTUD5
PFDN2
PPP1CA
SON
HEXIM1
STIP1
MIEF1
DPY30
FUT11

I

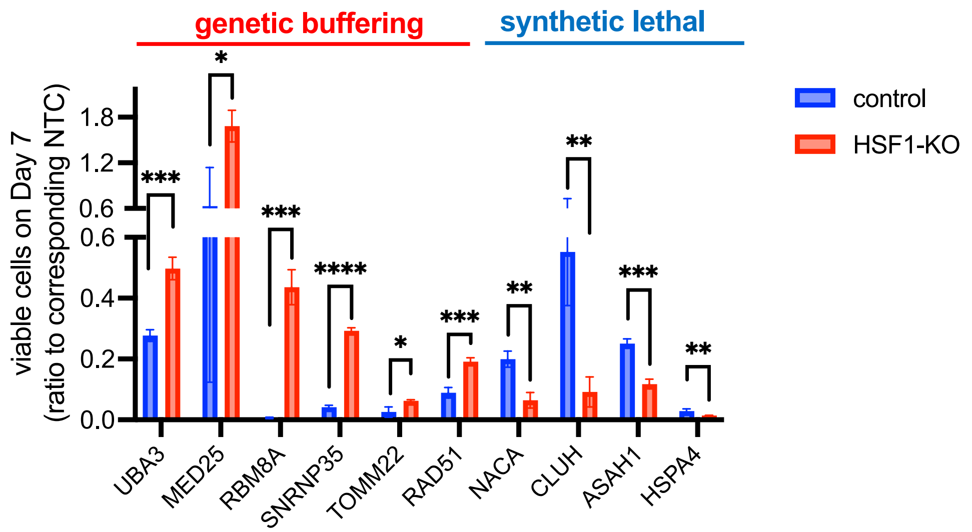
genetic buffering
synthetic lethal
viable cells on Day 7
(ratio to corresponding NTC)
1.8
1.2
0.6
0.6
0.4
0.2
0.0
control
HSF1-KO
UBA3
MED25
RBM8A
SNRNP35
TOMM22
RAD51
NACA
CLUH
ASAH1
HSPA4

Fig.S2_Nguyen et al.

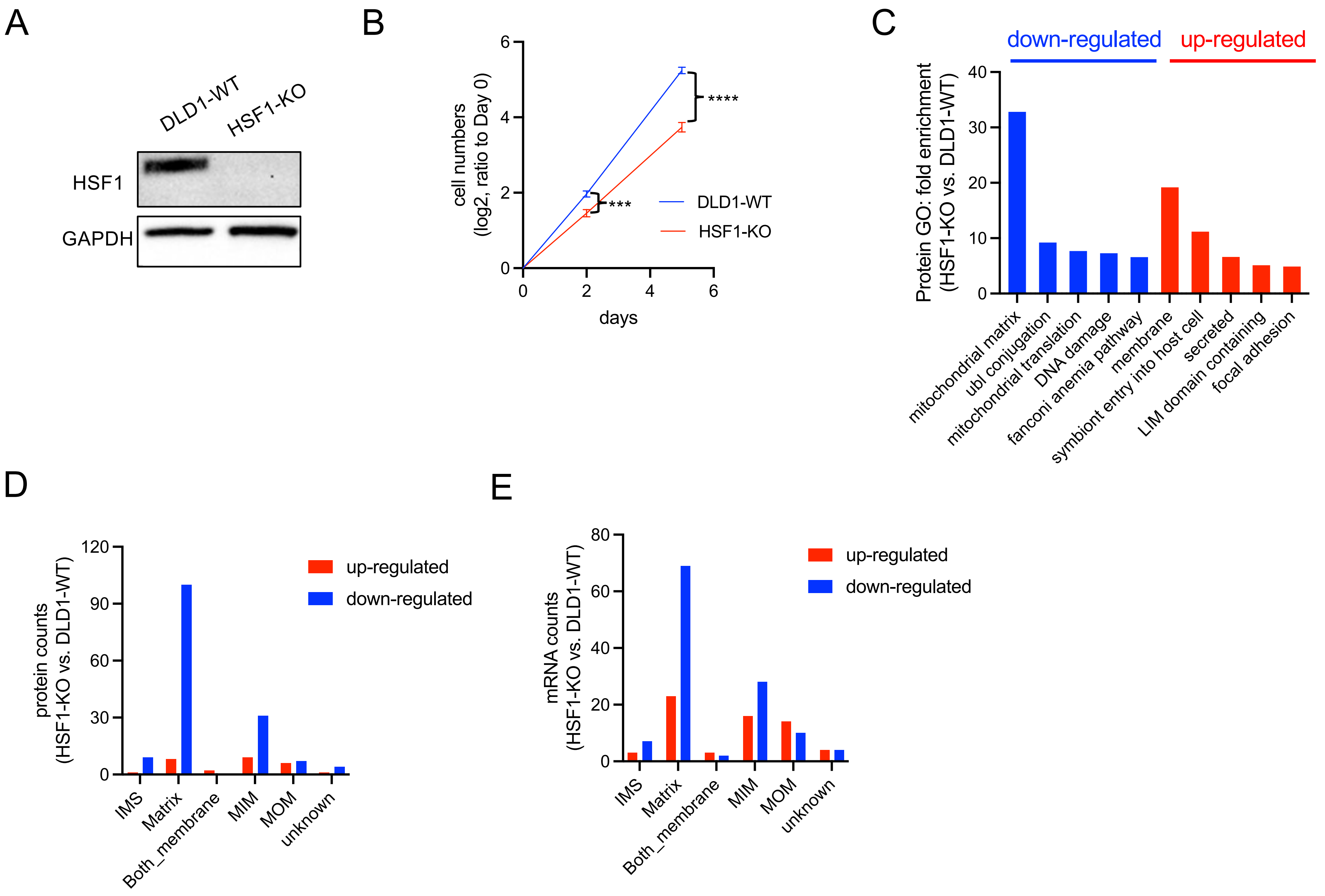

Fig.S3_Nguyen et al.

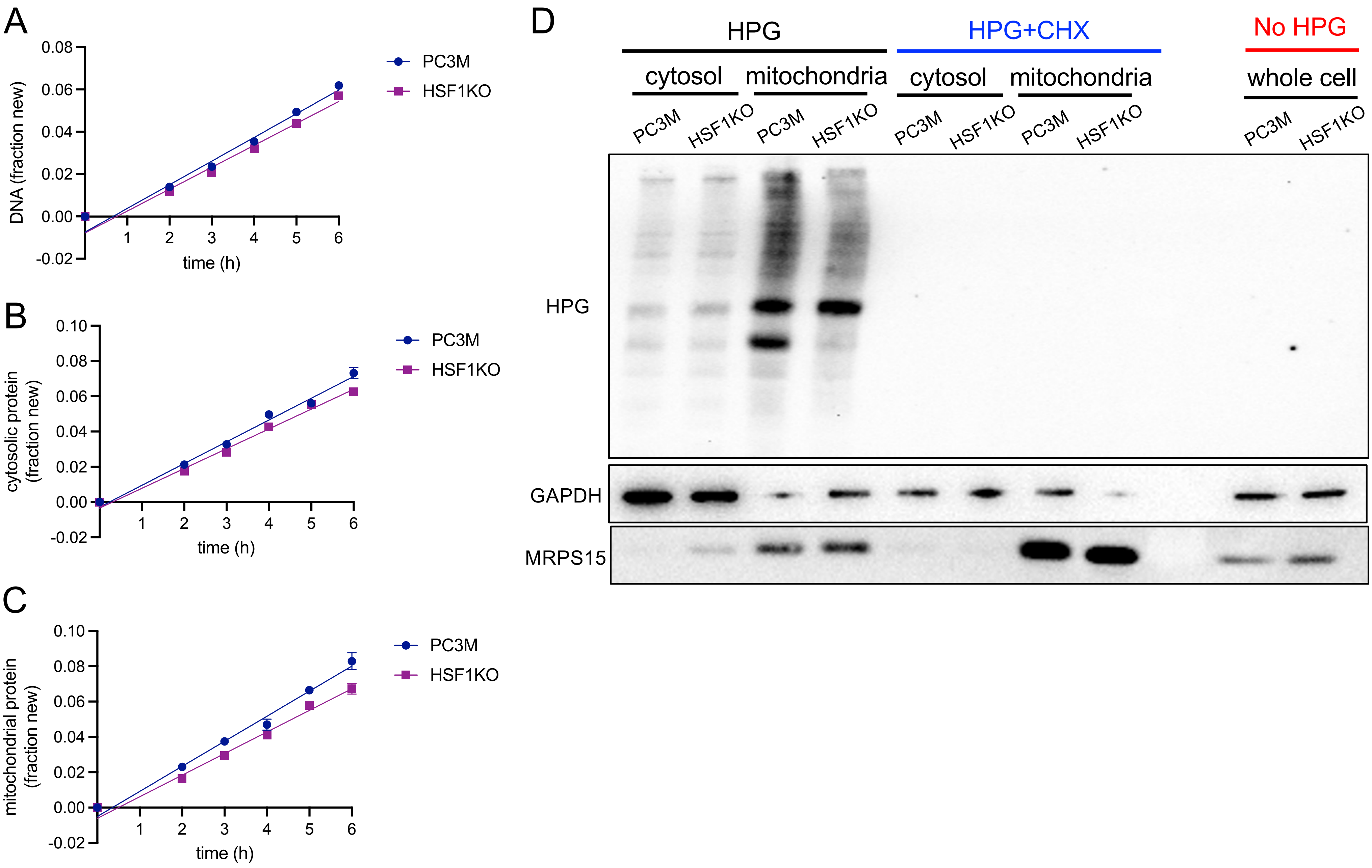

Fig.S4_Nguyen et al.

A

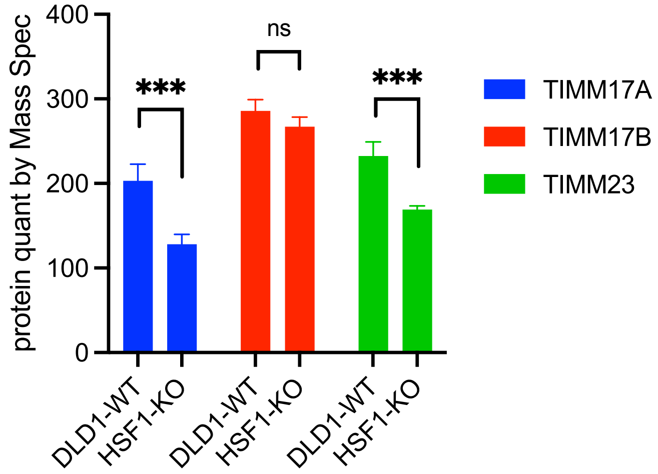


B

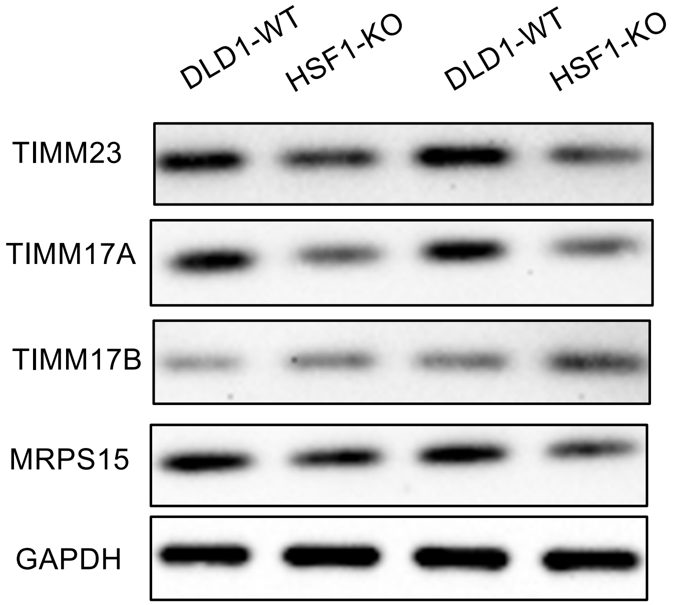

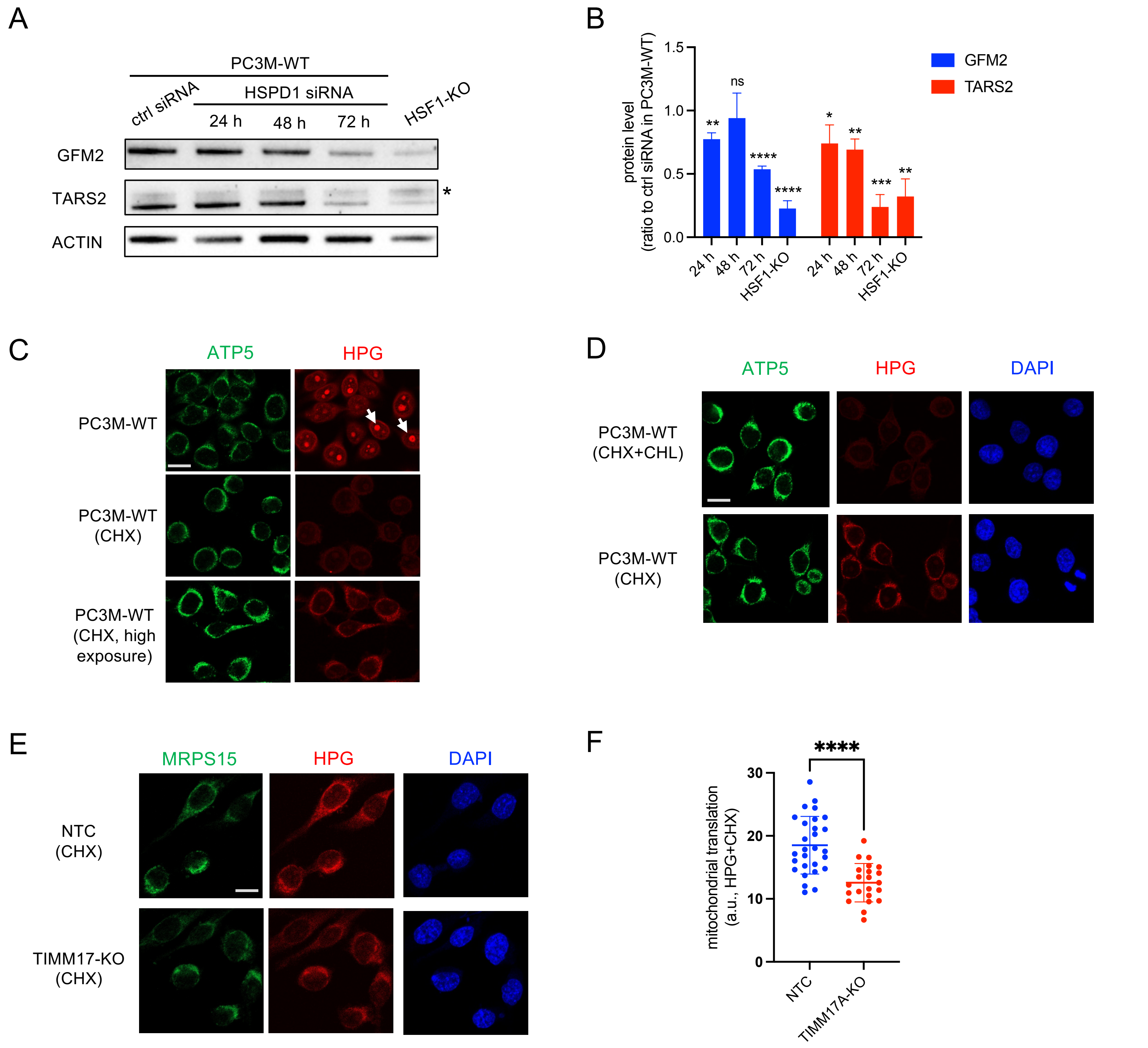
A
PC3M-WT
ctrl siRNA
HSPD1 siRNA
24 h
48 h
72 h
HSF1-KO
GFM2
TARS2
ACTIN
*
B
protein level
(ratio to ctrl siRNA in PC3M-WT)
1.5
1.0
0.5
0.0
**
ns
****
****
*
**
***
**
GFM2
TARS2
24 h
48 h
72 h
HSF1-KO
C
ATP5
HPG
PC3M-WT
PC3M-WT
(CHX)
PC3M-WT
(CHX, high
exposure)
D
ATP5
HPG
DAPI
PC3M-WT
(CHX+CHL)
PC3M-WT
(CHX)
E
MRPS15
HPG
DAPI
NTC
(CHX)
TIMM17-KO
(CHX)
F
****
mitochondrial translation
(a.u., HPG+CHX)
30
20
10
0
NTC
TIMM17A-KO

Fig.S6_Nguyen et al.

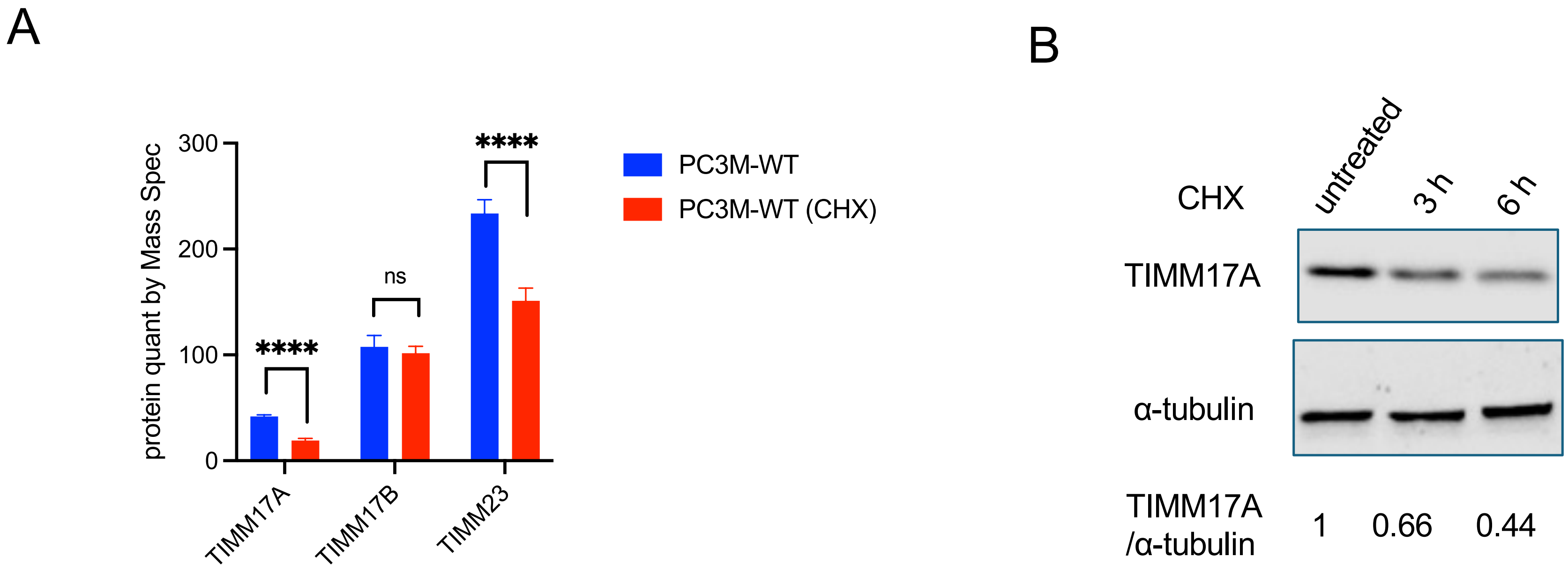

### Figure Legend

**Fig. S1, related to Fig. 1. HSF1 interactors are involved in critical cellular processes.**
(A) Growth curve of PC3M wild-type cells (PC3M-WT) and PC3M cells with HSF1 knockout (HSF1-KO). Cell numbers were counted every 48 h, and the ratios to the seeding cell numbers on Day 0 were converted to log2 fold change (mean ± SD, n=3). Paired t-test: * $P < 0.05$. The doubling time (mean ± SD, n=3): PC3M-WT: 26.1 ± 2.9 h; HSF1-KO: 29.3 ± 2.4 h.
(B) Demonstration of CRISPR score calculation using RAD51 in PC3M-WT cells as an example. The relative abundance of four sgRNAs targeting RAD51 over time is presented, and the weighted average is fitted into an exponential decay model.
(C) The distribution of the delta CRISPR score of non-target control sgRNAs from the first pair of screens. Our sequencing detected 994 out of 1000 control sgRNAs in the Brunello library. The synthetic lethal and genetic buffering cut-offs were set at a z-score of two. Based on the number of non-target control sgRNAs that fell outside the cut-offs, the p-values were less than 0.025. A stricter cut-off (z-score of three) was implemented for the second pair of screens, where a longer negative selection was performed.
(D-F) Histograms showing the viable cells measured by PrestoBlue assays on Day 4 or Day 7 in the co-CRISPR assays conducted in the Li Lab. The ratios to the corresponding NTC in either the control or HSF1-KO cells are presented (mean ± SD, n=3). Genes exhibiting a significant difference ($P<0.05$) between single KO and double KO (together with HSF1) on Day 4 (D&E), or on Day 7 but not on Day 4 (F), are shown (mean ± SD, n=3). Unpaired t-test: $P < 0.05$; ** $P < 0.01$; *** $P < 0.001$; **** $P < 0.0001$. Non-target control (NTC) and HSF1 serve as negative and positive controls, respectively. CHAF1B, which already displays a significant difference on Day 4, was included in the Day 7 test as a positive control. This set of co-CRISPR assays encompassed all 115 candidate HSF1 interactors from our screens (Table S1).
(G-I) Histograms displaying the viable cells measured by PrestoBlue assays on Day 4 or Day 7 in the co-CRISPR assays conducted in the Mendillo Lab. The ratios to the corresponding NTC in either the control or HSF1-KO cells are reported (mean ± SD, n=3). Genes that showed a significant difference ($P < 0.05$) between single KO and double KO (along with HSF1) on Day 4 (G&H), or those that did not show a difference on Day 4 but did on Day 7 (F), are shown (mean ± SD, n=3). Unpaired t-test: $P < 0.05$; ** $P < 0.01$; *** $P < 0.001$; **** $P < 0.0001$. This set of co-CRISPR assays included 110 of the 115 candidate HSF1 interactors from our screens, with the clones of SSRP1, SUPT4H1, IQGAP3, PLA2G12B, and CCAR1 that failed in infection.

**Fig. S2, related to Fig. 2. HSF1 remodels the steady-state mitochondrial proteome.**

(A) Western blot of DLD1 wild-type cells (DLD1-WT) and DLD1 cells with HSF1 knockout (HSF1-KO). GAPDH serves as a loading control.

(B) Growth curve of DLD1 wild-type cells (DLD1-WT) and DLD1 cells with HSF1 knockout (HSF1-KO). Cell numbers were counted every 48 h, and the ratios to the seeding cell numbers on Day 0 were converted to log2 fold change (mean $\pm$ SD, n=3). Paired t-test: *** P < 0.001; **** P<0.0001. The doubling time (mean $\pm$ SD, n=3): DLD1-WT: 23.08 $\pm$ 0.39 h, HSF1KO: 32.59 $\pm$ 1.11h.

(C) Histograms showing gene ontology (GO) analysis of differentially expressed proteins following the knockout of HSF1 in DLD1 cells (1.5-fold, FDR: 0.05).

(D&E) Histograms showing the numbers of differentially expressed proteins following HSF1 knockout in DLD1 cells across various mitochondrial compartments. Mitochondrial genes that significantly altered their protein (D) or mRNA (E) levels (1.5-fold, FDR: 0.05) were included. IMS: intermembrane space; MIM: mitochondrial inner membrane; MOM: mitochondrial outer membrane.

**Fig. S3, related to Fig. 3. HSF1 promotes mitochondrial biogenesis and function.**

(A-C) Scatter plots of newly synthesized DNA (A), cytosolic proteins (B), and mitochondrial proteins (C) as detected by deuterium oxide ($D_2O$) labeling in PC3M-WT and HSF1-KO cells (mean $\pm$ SD, n=3). Samples were collected hourly between 2 hours and 6 hours of labeling.

(D) Western blot analysis of newly synthesized cytosolic and mitochondrial proteins after one hour of HPG labeling in PC3M-WT and HSF1-KO cells. Cycloheximide (CHX) was added to a subset of labeling reactions to inhibit cytosolic translation. GAPDH and MPARS15 are used as cytosolic and mitochondrial protein markers, respectively.

**Fig. S4, related to Fig.4. HSF1 alters TIM complex composition.**

(A) Protein levels of the core TIM23 complex determined by TMT proteomics in DLD1 wild-type cells (DLD1-WT) and DLD1 cells with HSF1 knockout (HSF1-KO). Unpaired t-test: ns P >= 0.05; *** P < 0.001.

(B) Western blot analysis of the core TIM23 complex in biological duplicates. MRPS15 and GAPDH are loading controls for mitochondrial and total proteins.

**Fig. S5, related to Fig.6. TIMM17A promotes mitochondrial translation.**

(A&B) Representative images (A) and quantification (mean $\pm$ SD, n=3) (B) of western blot analysis of GFM2 and TARS2 upon HSPD1 siRNA treatment. In each experiment, the tested proteins were first normalized to the ACTIN loading control, and the ratios relative to the protein level in the control siRNA (ctrl) were calculated and plotted into histograms. Unpaired t-test (against control siRNA): ns $P >= 0.05$; * $P < 0.05$; ** $P < 0.01$; *** $P < 0.001$; **** $P < 0.0001$.

(C) Representative images of HPG labeling for newly synthesized proteins. HPG labeling was performed in wild-type PC3M cells (PC3M-WT) in the presence or absence of cycloheximide (CHX), which inhibits cytosolic translation. Arrowheads indicate nucleoli that show strong HPG signals. Immunofluorescence of ATP5 was used as a mitochondrial marker. Scare bar: 20 μM.

(D) Representative images of HPG labeling for newly synthesized proteins by mitochondrial translation. HPG labeling was performed in wild-type PC3M cells (PC3M-WT) in the presence of cycloheximide (CHX) with or without chloramphenicol (CHL), an inhibitor of mitochondrial translation. Immunofluorescence of ATP5 was used as a mitochondrial marker, and DAPI was used to stain DNA. Scare bar: 20 μM.

(E&F) Representative images (E) and quantification (mean ± SD) (F) of HPG labeling for newly synthesized proteins by mitochondrial translation. HPG labeling was performed in PC3M cells with TIMM17 knockout (TIMM17-KO) or the non-target control (NTC) for 45 min in L-methionine-free medium containing cycloheximide (CHX) to inhibit cytosolic translation. AAVS was used as the negative control in the co-CRISPR assay. Immunofluorescence of MRPS15 was used as a mitochondrial marker, and DAPI was used to stain DNA. Scare bar: 20 μM. Unpaired t-test: **** $P < 0.0001$.

**Fig. S6, related to Fig.7. TIMM17A protein level is coupled with HSF1 activity to promote robust cell proliferation.**

(A) Histograms showing the protein levels of TIM23 core complex components determined by TMT proteomics (n=4). The wildtype PC3M cells were treated with 100 μM of cycloheximide (CHX) for 6 hours or left untreated as the control. Unpaired t-test: ns $P >= 0.05$; **** $P < 0.0001$.

(B) Western blot analysis of TIMM17A in PC3M cells treated with 100 μM of cycloheximide (CHX) for 3 or 6 hours. TIMM17 protein was normalized to the α-tubulin loading control, and the ratios relative to that in the untreated control are labeled.
